# Small-molecule consumption drives metabolic stress and restricts erythroblast expansion in high-density cultures

**DOI:** 10.64898/2026.06.02.729469

**Authors:** Joan Sebastián Gallego-Murillo, Ingeborg van Lakwijk, Nurcan Yagci, Julie A. Reisz, Victoria Pozo Garcia, Angelo D’Alessandro, Luuk A. M. van der Wielen, Marieke von Lindern, Sebastian Aljoscha Wahl, Emile van den Akker

## Abstract

Transfusion-ready red blood cells can be cultured *ex vivo* from hematopoietic progenitors. Despite its promising outlook, a cultured transfusion unit cannot be produced at competitive costs. Large media volumes are required to maintain a maximum erythroblast cell density of 1-2.10^6^ cells/mL during the erythroblast proliferation stage. To identify the origin of the cell density limitation, we investigated the cellular support and metabolomic phenotype using different media formulations and feeding regimens. Media that were exposed to an increasing density of erythroblasts (termed spent media) displayed a proportional decrease in erythroblast proliferation support. A 1:1 combination of spent media with fresh media (not previously exposed to the cells) restored growth for all tested conditions. Filtering both fresh and spent media with a 3 kDa cut-off filter, and subsequent recombination of the two fractions, indicated that exhaustion of the small molecular weight fraction (<3 kDa) was primarily responsible for growth limitation. We performed targeted and untargeted metabolomics analysis, for both the intra- and extracellular compartments, following seeding in fresh medium (12, 24, 36 h). We observed degradation of nucleosides, depletion of amino acids, and a decrease in intermediates of the glutathione-ascorbate, γ-glutamyl and cysteine-methionine cycles. The latter compounds suggested an increase in oxidative stress in high density erythroblast cultures. Elimination of nucleosides from the medium led to a lower accumulation of purine salvage intermediates, and a 30% increase in cell productivity. In conclusion, we demonstrate that high-density erythroid cultures are subject to metabolic stress, defining critical constraints for scalable culture expansion.

## Introduction

Production of cultured RBCs (cRBC) has great potential as an additional source of transfusion-ready RBC, particularly to increase the availability of RBC with rare blood group phenotypes ^1–3^. Starting from induced pluripotent stem cells (iPSC)- or blood-derived hematopoietic stem cells, erythroblasts can be produced in large numbers ^4–6^. Expansion of erythroblast cultures requires the growth factors erythropoietin (Epo) and stem cell factor (SCF), as well as glucocorticoid hormones. Substitution of SCF and glucocorticoids by omniplasma, together with an increase in Epo levels, induces synchronous differentiation to mature, hemoglobinized, enucleated cRBC that resemble peripheral blood reticulocytes ^7–11^.

A single transfusion unit contains 2×10^12^ mature RBCs. To reach this high number of cells, a robust increase in cell numbers is needed during the expansion phase of the cultures. Currently, erythroblast cultures are kept at low cell densities (< 2×10^6^ cells/mL) through daily dilutions with fresh medium. Addition of fresh culture medium lowers the cell concentration, but also replenishes nutrients and growth factors, and dilutes potential inhibitory waste products and cytokines produced by the proliferating cells ^12–14^.

Growth limitations occur in high cell density erythroblast cultures (3-5×10^6^ cells/mL) after less than 20h of inoculation in fresh medium, suggesting either a quick depletion of essential nutrients or growth factors, and/or the fast accumulation of inhibitory cytokines and/or metabolic byproducts ^15^. Supplementation of dense cultures with specific nutrients, such as glucose, amino acids, vitamins, and/or with growth factors (Epo, SCF) does not restore the proliferation rate ^15^.

Growth inhibition is commonly observed in batch and fed-batch cultures of industrially relevant animal cell lines. Accumulation of lactate and ammonia is the most common cause of growth inhibition ^16^. Under substrate-excess conditions, rapidly proliferating cell cultures heavily rely on glycolysis, accumulating lactate, which may result in toxic effects due to increased osmolarity and decreased pH. Ammonia, mainly produced as a byproduct of glutamine catabolism, is also a common cause of growth inhibition ^17–21^. However, accumulation of lactate and ammonia in erythroblast cultures is well below growth inhibitory levels ^15,22–26^. Understanding the factors that influence erythroblast growth is crucial to upscale cultures for clinical applications.

Untargeted metabolomics in mammalian cell cultures is an unbiased approach to identify the metabolic phenotypes associated with growth inhibition ^27^. For instance, growth-inhibitory intermediates of amino acid catabolism appeared to impair oxidative stress control in Chinese hamster ovary (CHO) cell cultures ^28,29^. Reduced production of these inhibitors by metabolic engineering and modifications in the medium composition significantly improved CHO cell yields, as well as the increase in quality of the final product, typically monoclonal antibodies ^30–32^. A similar iterative approach to the industrial-scale production of erythroblasts is underway in the current scientific landscape. Mathematical models were previously used in erythroblast cultures to optimize medium refreshment frequency, reducing medium requirements in repeated batch culture strategy ^33^. However, the requirement of frequent fresh medium addition, combined with the high cost of media, still leads to non-competitive high production costs ^34,35^. To achieve cost-effective production of cRBC, the culture conditions must be optimized to allow high cell density cultures. In the present work, we aim to identify the molecular origins of the observed limitations of cell concentrations in erythroblast cultures. Such knowledge will enable the development of novel strategies to surpass current limitations while optimizing feeding regime and media composition.

## Results

### Limited production of cultured erythroblasts is independent of growth factor availability

To determine the cell proliferation capacity of the culture (Cellquin) medium, erythroblasts obtained from peripheral blood mononuclear cells (PBMC) of two independent donors were inoculated in fresh medium at three different cell densities (1, 3, and 5×10^6^ cells/mL). The erythroblasts were cultured for 48 h without medium addition or refreshment. All seeding conditions resulted in lower proliferation levels compared to those predicted by an exponential growth model fitted to the first 12 h of growth (Figure 1A). The deviation from exponential growth was larger and took place earlier for high cell density cultures (Figure 1B). The doubling time calculated from the growth rate fitted to the first 12 h of culture ranged from 17.4 h for cultures seeded at 1.3×10^6^ cells/mL, 31 h for cultures seeded at 3.6 ×10^6^ cells/mL, to 34.7 h for cultures seeded at 5.2×10^6^ cells/mL (Figure 1C). This indicates that medium became rate limiting at increased cell density within 12 h after start of the culture.

**Figure 1.**
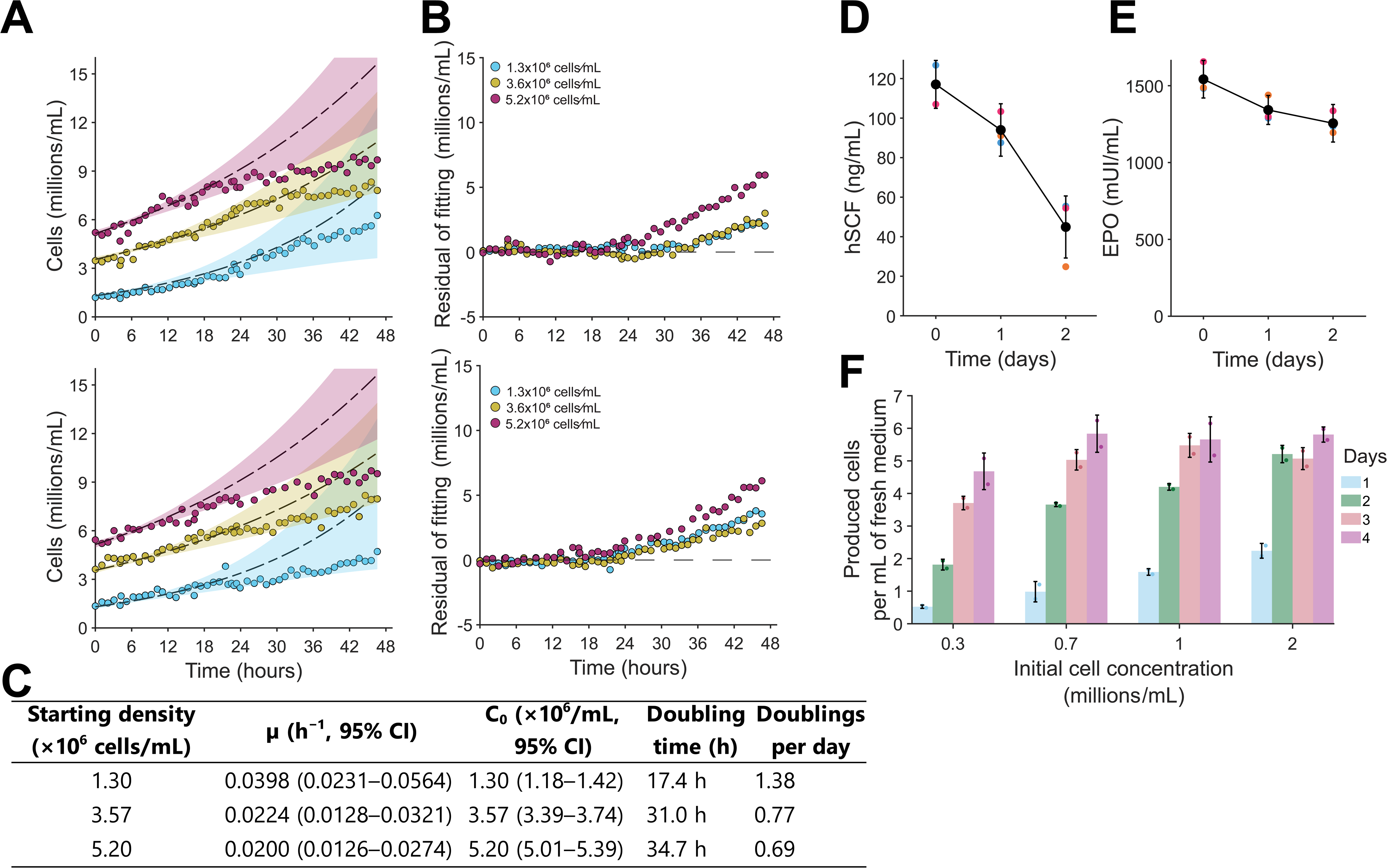
Growth limitations and maximal cell productivity in cultured proliferating erythroblasts. **(A-B)** PBMC-derived day 10 erythroblasts from two independent donors were inoculated in fresh proliferation medium at different starting cell densities (1.3, 3.6 and 5.2×10^6^ cells/mL), and cultured for 2 days without addition of fresh medium (batch culture), with hourly measurements of cell concentration. An exponential growth model, shown as black dotted lines, was fitted for each starting cell density to the first 12 hours of cell culture of both donors. Shaded areas indicate the upper and lower 95% prediction bounds for the fitted model (determined with MATLAB *predint* function) **(A)**. The difference between measured cell concentration and predictions of the exponential growth model was calculated through the complete culture time (**B**; colors match panel A). **(C)** Fitted parameters of exponential growth mode to erythroblast cultures at different starting cell concentrations. Growth rate (µ) and initial cell density (C_0_) of an exponential growth model were fitted to the first 12 hours of cell concentration measurements. Parameter values that minimized the sum of square errors are reported, together with 95% confidence bounds. Doubling time and doublings per day were calculated from the fitted growth rates. **(D-E)** Depletion levels of hSCF and Epo were determined in batch erythroblast cultures (3 independent donors) inoculated with a starting cell density of 1.0-1.5×10^6^ cells/mL. Colors indicate independent cultures, the black line indicates the average. **(F)** Maximum cell productivity (produced number of cells per mL of culture, calculated as the difference in cell concentration relative to the start of the culture) was determined for 4-day batch cultures inoculated at 0.3, 0.7, 1 and 2×10^6^ cells/mL using erythroblasts derived from 2 independent donors (growth curves available in Supplementary Figure 1). Error bars are displayed as standard deviations of measurements.

To determine whether depletion of the growth factors Epo or hSCF could explain the observed growth limitations, erythroblasts from three different donors were seeded at 1×10^6^ cells/mL in fresh medium supplemented with hSCF and Epo, and cultured for 2 days without medium refreshment. An average of 25% Epo and 50-80% of the initial hSCF were depleted at day 2 (Figure 1D-E). To determine whether periodical replenishment of growth factors would improve cell growth, erythroblasts seeded at cell concentrations between 0.3 and 2×10^6^ cells/mL were kept in culture for 4 days with daily addition of fresh hSCF and Epo (100 ng/mL and 2 IU/mL, respectively). After 2 days, growth in low cell density cultures slowed down, where high cell density cultures already reached a cell density plateau after 2 days (Supplementary Figure S1). Total number of cells produced per mL of medium relative to the start of the culture peaked at 5-6×10^6^ cells/mL for inoculation densities of 1-2×10^6^ cells/mL (Figure 1F; growth curves in Supplementary Figure S1). Thus, daily addition of growth factors did not alleviate the growth restrictions at high cell density.

The results suggest that growth factor depletion is not the origin of the growth decrease in batch cultures. Rather, there is a growth-linked depletion of one or more essential nutrients or the accumulation of toxic byproducts during the batch culture, as is often observed for other bioprocesses ^36^. Other medium feeding strategies are required for erythroblast cultures such as perfusion to increase nutrient supply and removal of inhibitory metabolites accumulated during culture.

### Perfusion bioreactor increases erythroblast cell density

Oxygen may be limiting in static erythroid cultures in dishes ^26^. However, limitations in cell productivity in batch erythroblast cultures are similar to erythroid stirred tank bioreactor cultures, in which oxygen is sufficiently provided^15^, suggesting that nutrient availability may be limiting. To determine which cultivation strategies can ensure sufficient nutrient availability, and decrease toxic byproducts, we modeled erythroblast proliferation bioreactor cultures under different feeding regimes: (1) batch (no feeding regime), (2) fed batch with constant feed rate (FB_const_), (3) fed batch with exponentially increasing feeding (FB_exp_), (4) perfusion with a constant perfusion rate (PERF_const_), and (5) perfusion at a fixed cell-specific perfusion rate (PERF_CSPR_) (Figure 2A).

**Figure 2.**
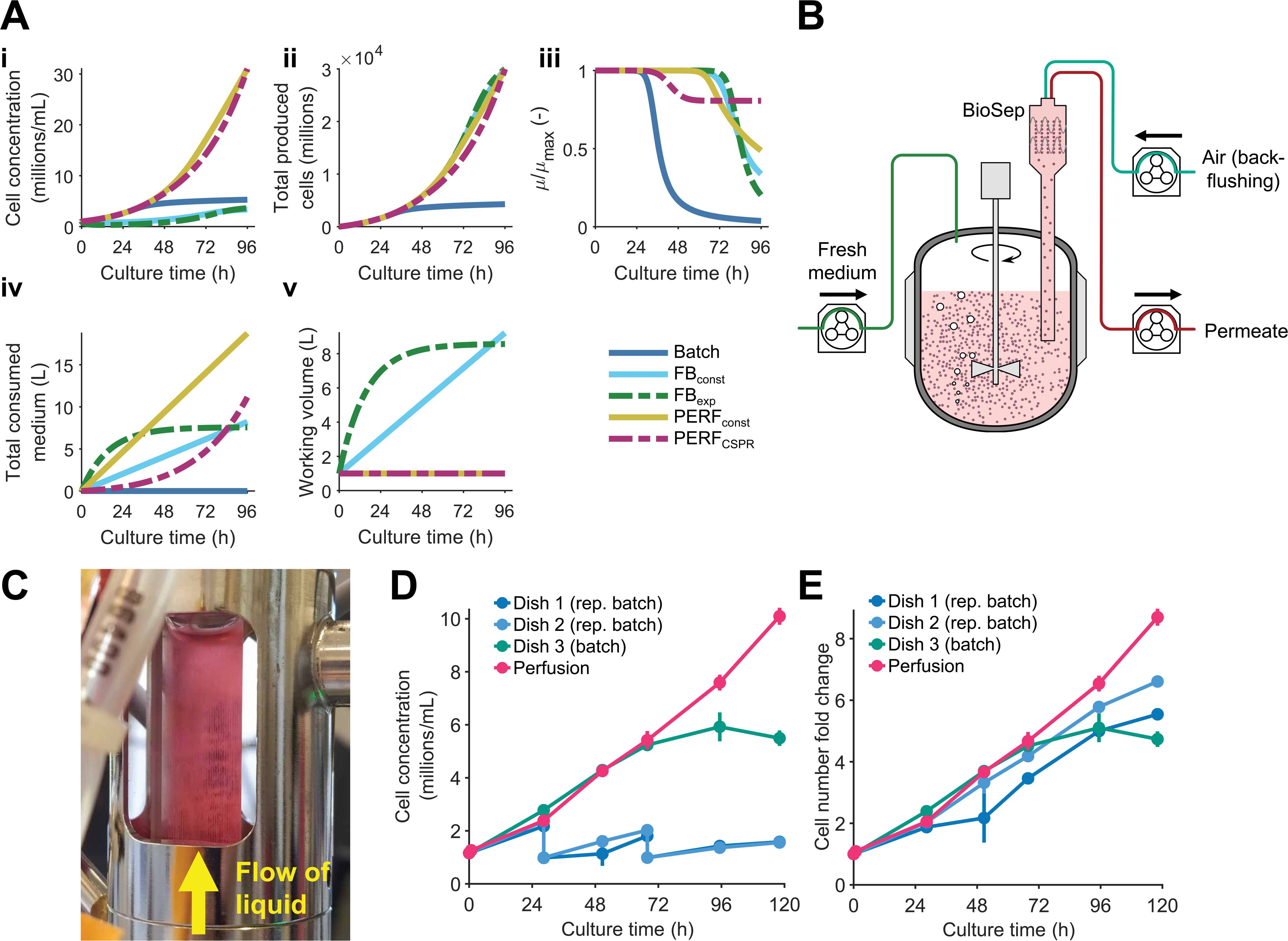
Perfusion allows to surpass cell-density limitations in erythroblast proliferation cultures. **(A)** Bioreactor cultures with different feeding strategies (batch, fed batch with constant feed rate *FB_const_*, fed batch with exponentially increasing feed *FB_exp_*, perfusion at a constant rate *PERF_const_*, perfusion at a constant cell-specific perfusion rate *PERF_CSPR_*) were simulated, using a kinetic growth model for erythroblast proliferation including a putative inhibitor produced in a growth-dependent manner. Except for batch mode, the feeding parameters of each feeding regime were optimized to achieve a production of 30×10^9^ cells in 4 days in a 1 L bioreactor inoculated at a starting cell concentration of 1×10^6^ cells/mL, minimizing the medium volume requirements. Cell concentration **(Ai)**, total produced cells **(Aii)**, magnitude of the growth inhibition (calculated as the ratio of the instantaneous growth rate µ and the maximum growth rate µmax; **Aiii)**, total volume of medium consumed **(Aiv)**, and reactor working volume **(Av)** are displayed. Equations used for the growth model and for the bioreactor simulations are available in Supplementary Methods. **(B)** Day 9 erythroblasts were cultured in a 0.5 L stirred tank bioreactor (working volume = 250 mL) with perfusion, using an acoustic cell retention device (Applikon BioSep), operated semi-continuously, with cycles of permeate liquid flow and cell backflushing using air. **(C)** In this system, cells are retained in the cuvette by sound waves against the upward movement of liquid. Cells can be seen as vertical lines in the separation chamber. **(D-E)** Erythroblast concentration (D) and fold-change increase of erythroblast numbers (E) in a 5 day culture in a stirred tank bioreactor using an average cell-specific perfusion rate of 500 pL/cell/day (red line), and in culture dishes following either a batch strategy without feeding (green line) or a repeated batch strategy in which the erythroblast concentration was restricted to <2×10^6^ by dilution with fresh medium (blue lines). Perfusion rates during the complete bioreactor run are available in Supplementary Figure S3.

For these simulations, we used the kinetic model previously reported by Glen et al., in which erythroblast growth limitation, either via media exhaustion or toxification, was dependent on cell growth ^37^. This model also included an inhibition decay mechanism (model details available in Supplementary Methods).

A 1 L culture with 1×10^6^ cells/mL was used as starting condition for all scenarios, and the optimized feeding parameters for each culture strategy were determined to minimize the overall medium consumption while reaching a total yield of 30×10^9^ cells after 4 days of culture. All evaluated perfusion feeding regimes led to higher cell densities (Figure 2Ai) and yield (Figure 2Aii) compared to conventional batch culture mode. Both fed-batch alternatives (FB_const_, FB_exp_) and CSPR-based perfusion (PERF_CSPR_) led to similar medium consumption levels (Figure 2Aiii), but the medium additions used in fed-batch cultures require a bioreactor working volume of 8 L at the end of the culture (Figure 2Aiv). For FB_exp_, a negative exponential factor was found to be optimal, equivalent to fast feeding in the first days of culture, followed by slow feeding at the end of the cultivation. Under constant CSPR perfusion, maximal inhibition in cell growth rate was <20% during the 4 days of culture, while for fed-batch the growth was reduced by >50% (Figure 2Aiii). Similar values for the optimized parameters of each feeding regime were obtained for both evaluated models (Supplementary Figure S2, Supplementary Table S1).

To validate whether CSPR-based perfusion allows for higher cell concentrations under our own cell culture conditions, day 9 erythroblast cultures were seeded in a 500 mL STR (working volume = 250 mL) at an initial cell concentration of 1×10^6^ cells/mL. Fresh medium was continuously perfused into the bioreactor, and cells were retained using an acoustic filtration system (Figure 2B-C). An average daily CSPR of 500 pL/cell/day was used (daily perfusion rates available in Supplementary Figure S3), equivalent to the medium requirements of culture dishes seeded at the same initial concentration and with half medium refreshment every day. After 5 days of culture, cultures in the perfused bioreactor reached a cell concentration of 1×10^7^ cells/mL (Figure 2D), with similar overall cell number fold change per 24 h compared to static cultures in dishes under a repeated batch feeding regime (daily reseeding at 1×10^6^ cells/mL with fresh medium) (Figure 2E). A culture dish kept without medium addition (batch mode) showed a decrease in growth after 3 days of culture (Figure 2 D, E and Supplementary Figure S1).

Thus, experimental data confirmed increased erythroblast concentrations in a fixed culture volume that can be achieved by a perfusion feeding approach. The observations highlight that the cause of the growth limitation in erythroblast proliferation cultures can be mitigated by the supply of fresh nutrients and/or removal of toxic byproducts generated during growth.

### Exhaustion of small molecules impairs erythroblast proliferation

During perfusion, fresh growth factors and metabolites were fed to the reactor, while potential toxic metabolites were removed from the bioreactor by continuous medium exchange. We next determined whether growth decrease was caused by the depletion of an essential nutrient or the accumulation of a toxic byproduct in culture medium. For this, erythroblasts were seeded in fresh medium at concentrations between 1 and 15×10^6^ cells/mL overnight (16 h). The spent medium (after 16 h cell exposure) was filter sterilized, supplemented with fresh growth factors (Epo, hSCF, and dexamethasone), and used to culture an independent batch of erythroblasts at identical low cell densities (0.7×10^6^ cells/mL). When seeded in the spent medium obtained from various cell densities, erythroblasts doubled the first 24 h, but the cell yield progressively and significantly decreased in the next 24 h with spent medium derived from

increasing cell densities (Figure 3A). Spent media derived from 15x10^6^ cell/mL condition almost completely abolished proliferation (Figure 3A). Thus, the proliferation of fresh cells was increasingly inhibited when media were pre-exposed to higher erythroblast densities. Dilution of spent media with fresh medium (1:1) did not shift this correlation between proliferation of fresh cells and the number of cells used in pre-exposure though cell proliferation was recovered for all conditions (Figure 3B). This suggests that the decrease in cell proliferation at high cell density is caused by nutrient depletion rather than the production of a toxic byproduct, with the depleted nutrient(s) replenished by the addition of fresh medium.

**Figure 3.**
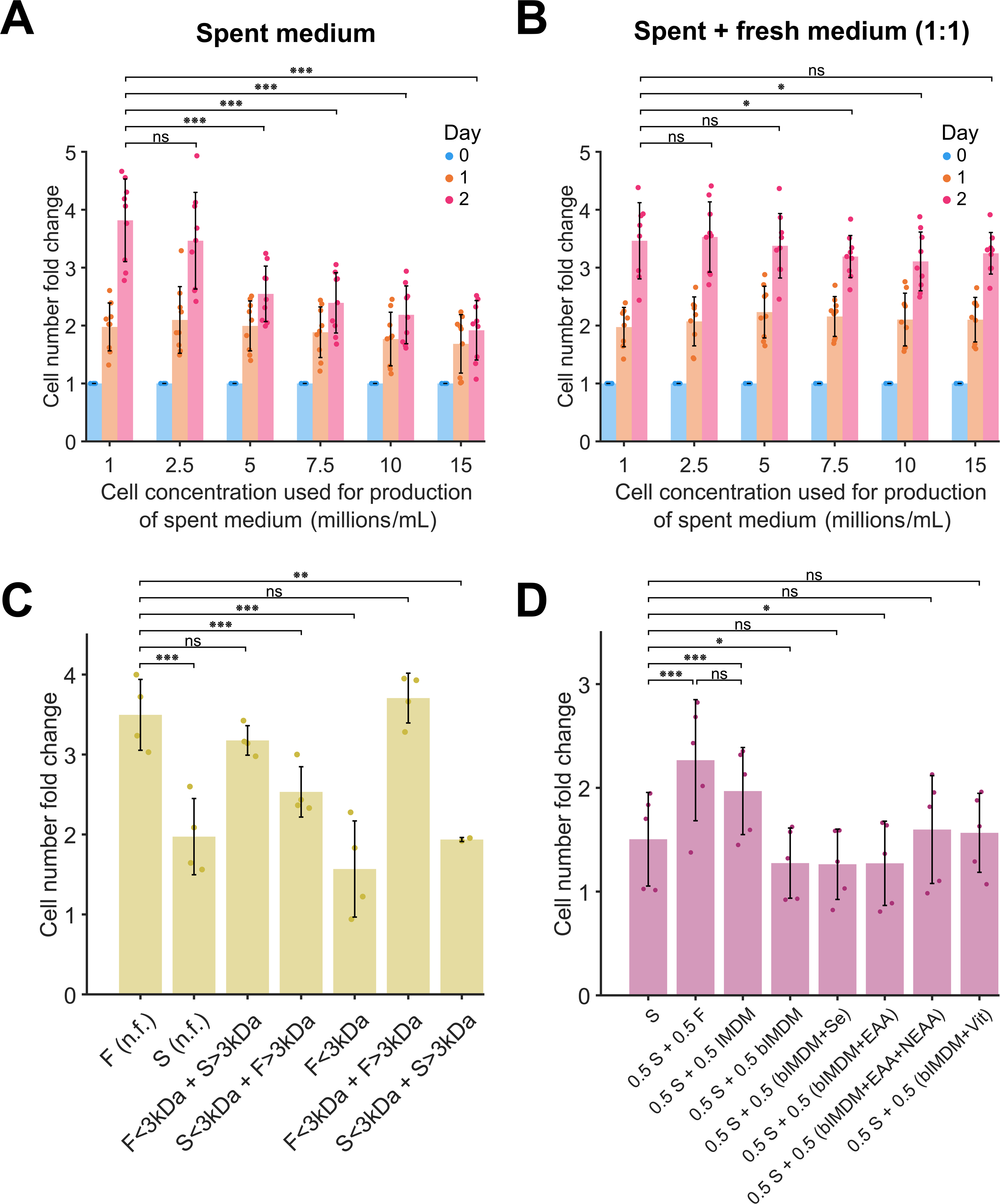
Small molecules (<3 kDa) contribute the most to the observed growth inhibition in erythroblast batch cultures. **(A-B)** Spent medium was generated culturing erythroblasts for 16h at different starting cell concentrations (1-15×10^6^ cells/mL). Either only spent medium **(A)** or a 1:1 mix of spent and fresh media **(B)** was used to seed fresh erythroblasts, which were cultured for 2 days without any media addition (batch cultivation). **(C)** Fresh (F) and spent (S) media were fractionated via ultrafiltration (size threshold: 3 kDa). Erythroblasts were inoculated in different combinations of the flowthrough (<3 kDa) and retentate (>3 kDa) of both media. Cell number fold change after 2 days of culture without refreshment in the reconstituted media is displayed. **(D)** Effect of supplementing spent medium with solutions containing specific components present in Iscove’s Modified Dulbecco’s Medium (IMDI) on erythroblast proliferation level. Basal IMDM (bIMDM), only containing glucose and the inorganic salts present in full IMDM, was used as background solution to which other IMDM components were added (sodium selenite, Se; essential amino acids, EAA; non-essential amino acids, NEAA; vitamins, Vit). Cell number fold change for erythroblasts inoculated after 2 days of culture is displayed. All data are displayed as mean ± SD (error bars; *n* = 9 donors for panels A and B, *n* = 4 for panel C, *n* = 4 for panel D). Significance levels were calculated using paired two-tailed Student’s t-test (ns for non-significant, * for *p* < 0.05, ** for *p* < 0.01, *** for *p* < 0.005).

Erythroblast culture media contains small molecules (e.g. glucose, amino acids, vitamins, trace elements), proteins (albumin, transferrin, insulin), and lipids. To determine which fraction of medium components is depleted, both fresh and spent medium (from 15×10^6^ cells/mL cultures) were filtered through a 3 kDa cutoff filter. The retentate (>3 kDa) and flow-through (<3 kDa) fractions from fresh (F) and spent (S) medium were recombined and used for cultures of erythroblasts inoculated at 0.7×10^6^ cells/mL with addition of fresh growth factors.

The control recombination of retentate and flow-through fractions of fresh medium resulted in similar cell proliferation compared to the unfractionated medium. Unfractionated and recombined spent medium also resulted in similarly reduced cell proliferation (Figure 3C). Importantly, cell growth was restored when erythroblasts were seeded in spent medium retentate combined with flow-through from fresh medium (F<3kDa + S>3kDa). In contrast, cell growth was reduced when erythroblasts were seeded in fresh medium retentate and spent medium flow-through (S<3kDa +F>3kDa). Furthermore, small molecules from fresh medium alone (F<3kDa only) were not sufficient to restore growth. These results suggest that exhaustion of small molecules <3kDa contributed to the growth limitation in erythroblast batch cultures.

To identify putative depleted small molecules, the spent medium was supplemented with key components of the basal medium (IMDM; used to generate Cellquin), including essential and non-essential amino acids (EAA and NEAA, respectively), vitamins, and sodium selenite (Figure 3D). Compared to spend medium condition, supplementing the culture with half fresh Cellquin (F) or IMDM significantly increased cell growth, albeit that recovery was somewhat lower for IMDM. No increase in growth was observed with the addition (to IMDM media) of non-essential amino acids and vitamins, and no single condition led to a full recovery of erythroblast proliferation to levels observed in fresh medium (Figure 3D). The results prompted us to investigate the kinetics of nutrient consumption.

### Metabolic profiles of erythroblast culture reveal amino acids to be heavily consumed during proliferation

To obtain a dynamic view of medium nutrients and intermediate metabolites during culture, metabolic profiling was performed for the extra- and intra-cellular compartments of the culture. Erythroblasts cultured from three independent donors were seeded in fresh medium at 0.7 and 2×10^6^ cells/mL and cultured for 12, 24, and 36 h. Both cell pellets and supernatant were analyzed by liquid chromatography coupled to mass spectrometry (LC-MS) metabolomics (Supplementary data 1). Data was acquired using untargeted acquisition followed by a targeted data analysis focusing on pathways central to energy and redox metabolism.

Analysis of supernatant samples identified 93 annotated metabolites. Differentially detected features were determined by comparison to blank samples run before and after the analysis, and metabolites that showed high variation (relative standard deviation RSD > 25%) in quality control samples were not considered for further analysis (Supplementary Figure S4). Principal component analysis (PCA) of extracellular metabolite data showed that independent biological replicates (donors) of a specific time point clustered together. The first two components described 88% of the total variance (PC1 = 75%, PC2 = 13%; Figure 4A). The clustering of samples appeared dependent on time and initial culture density. Central carbon metabolism intermediates and nucleosides largely contribute to PC1 (Figure 4B). Hypoxanthine, an intermediate in purine degradation, displayed the strongest loading (absolute value) for PC2. For a comprehensive overview of the common and discriminating profiles, metabolite changes were visualized using k-means clustering (k=5) (Figure 4C-D; see Supplementary Figure S5 for the determination of the optimal number of clusters). Most metabolites showed a monotonic decrease in extracellular concentration with culture time as shown in cluster 4 (yellow). This cluster was largely composed of amino acids, tryptophan catabolites, and arginine pathway metabolites. On the other hand, cluster 1 metabolites (red, e.g. hypoxanthine) displayed a transient accumulation while cluster 2 (orange) represents transiently depleted metabolites. Dicarboxylates of the tricarboxylic acid (TCA) cycle (e.g., citrate, 2-oxoglutarate, malate, and fumarate) accumulated with time as did lactate, acidic amino acids (e.g. Asp, Glu), and polyamines spermidine and spermine (cluster 5). In contrast, nucleoside and nucleobase concentrations rapidly decreased (e.g., guanosine and adenosine; cluster 3).

**Figure 4.**
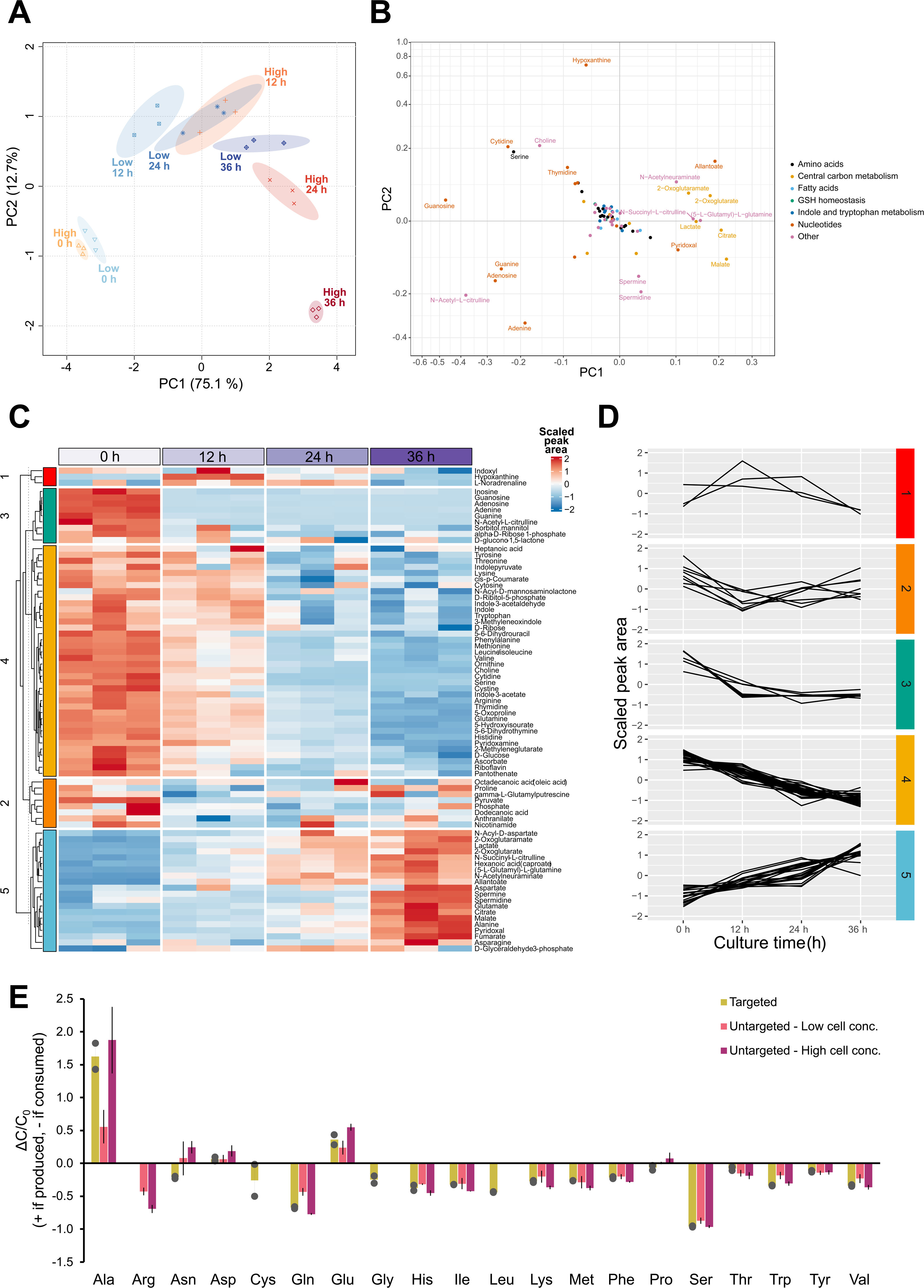
Metabolomics analysis of erythroblast batch culture supernatant. **(A-D)** Erythroblasts from three independent donors were expanded from PBMCs for 11 days, subsequently inoculated in fresh proliferation medium at a low (0.7×10^6^ cells/mL) or high (2.0×10^6^ cells/mL) cell concentration, and cultured without medium refreshment for 36 h. Supernatant samples taken at 0, 12, 24, and 36 h were analyzed by untargeted UHPLC-MS metabolomics. **(A)** After removal of non-informative features (background), and normalization (glog transformation + mean centering) of the peak area, principal component analysis (PCA) was performed. **(B)** Biplot with the loadings of all metabolites for the first two principal components (PC1, PC2). Only metabolites with loadings > 0.1 are labelled. **(C)** Heat map of metabolite peak areas for the high cell density cultures. **(D)** Metabolites were *k*-means clustered, resulting in five groups of metabolites displaying similar trends over the timeseries (cluster 1, *n*=3 metabolites; cluster 2, *n*=9; cluster 3, *n*=38; cluster 4. *n*=8; cluster 5, *n*=20). **(E)** Validation of the amino acid consumption levels observed in the untargeted metabolomics analysis was performed via quantitative determination of amino acid concentrations by GC-MS. Erythroblast from two donors were cultured in fresh medium for 2 days. Amino acid concentrations were measured at the beginning (*C_0_*) and end of the culture (*C*) (data available in Supplementary Table S2). Relative change in amino acid levels is calculated as Δ𝐶⁄𝐶_0_ = (𝐶 − 𝐶_0_)⁄𝐶_0_ (>0 if produced, <0 if consumed), using the measured concentration from targeted metabolomics, or raw peak area for the untargeted data. For the latter, no data is available for glycine nor cysteine (it could not be measured with the used analytical platform). Data in panel E is displayed as mean ± range of the measurements (error bars; *n* = 2).

The contrasting trends of groups of amino acids warranted further investigation. Amino acids are major contributors to protein synthesis and, by extension, proliferation ^38^. Analysis of the top changed supernatant amino acids as a function of time shows time-dependent decreases in most amino acids with strongest downward trends in higher density cell culture media. Of these, serine, a non-essential amino acid, displayed low extracellular levels after 36 h of culture, both for low and high cell density cultures and was the top changed amino acid (Figure 4E). Amino acid changes were validated by targeted gas chromatography coupled to MS (GC-MS) measurements using cells from two independent donors seeded at 1×10^6^ cells/mL and cultured for 2 days (final cell concentration = 3×10^6^ cells/mL; amino acid concentrations available in Supplementary Table S2). Glutamine, the most abundant amino acid in fresh medium, was rapidly consumed (overall cell-specific consumption rate of 283-314 nmol/million cells/day; Supplementary Table S3), reaching a consumption level of 67% at the end of culture.

### The intracellular erythroblast metabolome shows analogous results to supernatant analyses

Intracellular metabolomics data was collected in parallel to the medium samples. Similar to the extracellular metabolome, the principal component analysis shows dependence with culture time, although with larger dispersion between biological replicates (Figure 5A-B). The composition of the metabolome of cells cultured at low concentration for 12 or 24 h overlapped with erythroblasts inoculated at high density for 12 h both in PC1 and PC2. Also the subsequent samples (36 h low density and 24 h high density) show similarities. Cells cultured for 36 h at high density were distant from other measured timepoints. Using k-means clustering (k=5), the main profiles were identified. Clusters 1 and 2 represent metabolites that are rapidly depleted, including several amino acids (serine, arginine, lysine), arginine pathway metabolites (creatine, creatinine, putrescine, *N*-acetylcitrulline), metabolites downstream of glucose (glucose 6-phosphate), pentose phosphate pathway metabolites (sedoheptulose phosphate, glucono-1,5-lactone phosphate) and purines (adenine, adenosine, guanosine Figure 5C-D). Cluster 3 represents metabolites that stay constant for 24 h but display accelerated decrease in intracellular levels at 36 h. These include glucose, ribose, PPP metabolites (erythrose phosphate, 6-phosphogluconate, glucono-1,5-lactone), pyrimidines, the majority of the amino acids, and redox-centric metabolites like glutathione disulfide, S-glutathionylcysteine, cysteine, and cystine (i.e. cysteine disulfide). Clusters 4 and 5 contain metabolites that initially increase in concentration but are reduced again after 36 h in culture. These include key energy currency ATP along with other mono- and diphosphate nucleosides plus energy metabolites of the TCA cycle and glycolysis.

**Figure 5.**
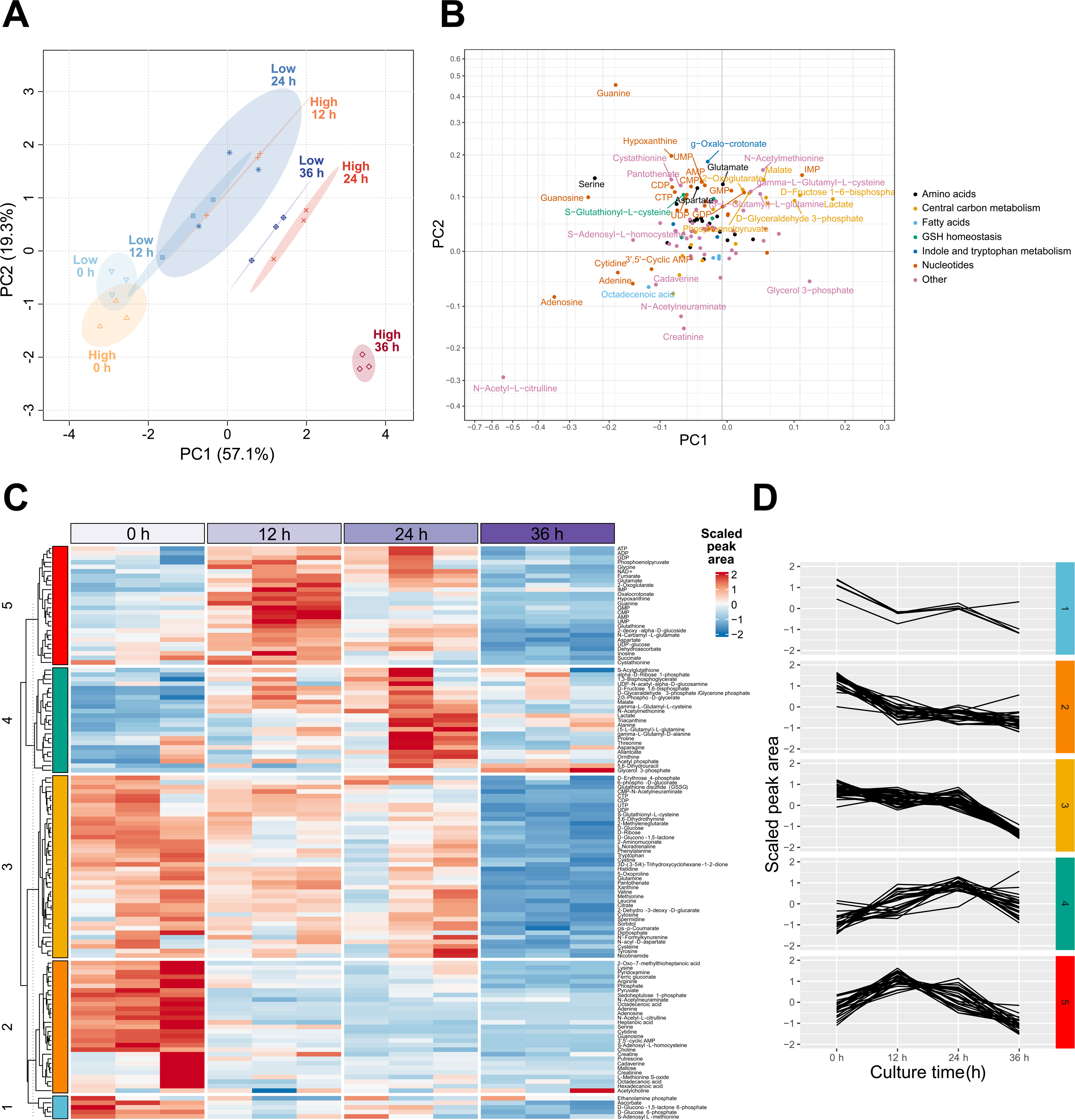
Statistical analysis of intracellular metabolomics in erythroblast proliferation cultures. Erythroblasts from three independent donors were expanded from PBMCs for 11 days, subsequently inoculated in fresh proliferation medium at a low (0.7×10^6^ cells/mL) or high (2.0×10^6^ cells/mL) cell concentration, and cultured without medium refreshment for 36 h. In parallel to the supernatant samples used for Figure 4, cell pellets were taken at 0, 12, 24, and 36 h. Intracellular metabolites were extracted using methanol and acetonitrile. Principal component analysis was performed using the normalized peak area data (**A**, score plot; **B**, loadings biplot). **(C)** Metabolites were clustered in 5 groups using *k*-means clustering performed over the high cell density culture peak area data (cluster 1, *n*=5 metabolites; cluster 2, *n*=29; cluster 3, *n*=40; cluster 4, *n*=23; cluster 5, *n*=26). **(D)** Metabolite peak area trends for each cluster.

The prominent depletion of amino acids both in medium and in cells suggests that this could be an inhibiting factor. Supplementation with amino acids, however, did not recover proliferation of erythroblasts in exhausted medium (Figure 3D). Thus, the depletion of amino acids likely contributes to the observed erythroblast growth limitations; nonetheless, other factors may play a role as well. The observed larger dispersion between samples in the PCA for intracellular data compared to extracellular data suggests that intracellular metabolite levels show less time-dependency compared to concentrations in the supernatant, indicating intracellular homeostasis.

### Nucleosides and deoxynucleosides in culture medium reduce cell growth

To better understand the implications of metabolites that increase or decrease over time, pathway-enrichment analysis was applied ^39–41^. Metabolite set quantitative enrichment analysis (MSQEA) was performed on the extracellular metabolome data of high cell density cultures (2.0×10^6^ cells/mL), which revealed an enrichment of purine metabolism in all timepoints relative to the start of the culture (Supplementary Figure S6). Purine metabolism comprises de novo biosynthesis from ribose 5-phosphate via phosphoribosyl pyrophosphate, purine salvage, and purine degradation ^42,43^. In cultured erythroblasts, net depletions of adenine, adenosine, guanosine, and inosine were observed intracellularly (Figure 6A), consistent with the decreases of these observed extracellularly (Figure 4C). This was accompanied by a transient accumulation of other intermediates of the purine degradation pathway such as hypoxanthine (Figure 6A). Hypoxanthine can be recycled to inosine monophosphate (IMP) or be further metabolized into xanthine and urate by xanthine oxidase, with superoxide radical anions and hydrogen preoxide as byproducts^44^. Although human cells lack the urate oxidase enzyme, accumulation of allantoate was observed extracellularly in the first 12 h of culture, while displaying a peak of intracellular levels after 24 h followed by a decrease in the following 12 h. The spontaneous oxidation of urate drives allantoate production, and this process produces 2 hydrogen peroxide molecules.

**Figure 6.**
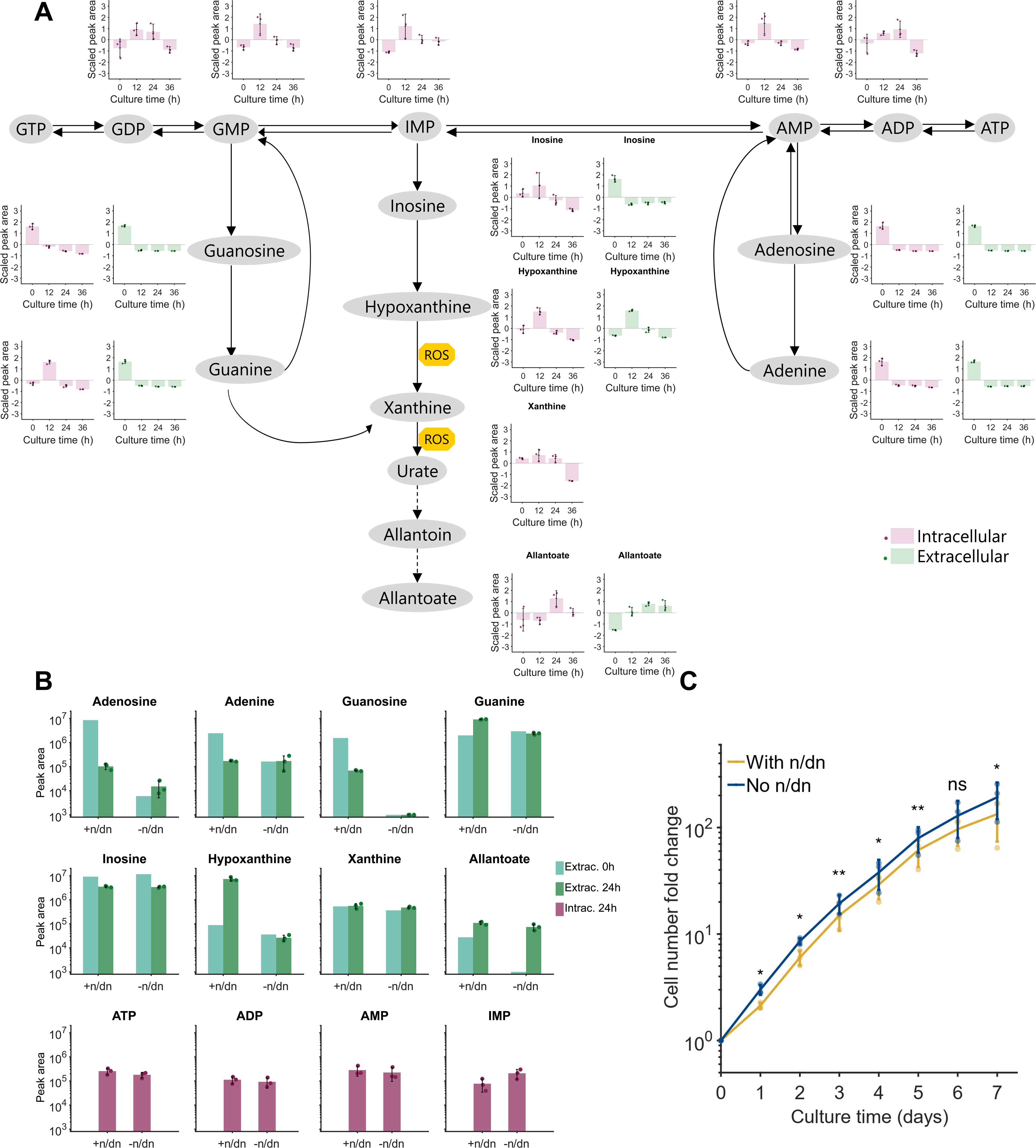
Cultured erythroblasts show high activity of the purine degradation pathway. **(A)** Extracellular (green) and intracellular (red) levels of purine degradation pathway intermediates for erythroblast (day 11) cultured at a starting cell concentration of 2.0×10^6^ cells/mL for 36 h in fresh proliferation medium. Although human cells do not have the enzymes required for the conversion of urate into allantoin and allantoate (dotted arrows), non-enzymatic processes can lead to those conversions. Some metabolites were only detected intra- or extracellularly. **(B)** To evaluate the effect of eliminating nucleosides (n) and deoxynucleosides (dn) from the medium formulation on the metabolism of cultured erythroblasts metabolism, cells were cultured for 24 h in medium with (+n/dn) and without (-n/dn) (deoxy-)nucleosides. Extracellular and intracellular levels of purine degradation pathway intermediates are displayed. **(C)** Medium without (deoxy-)nucleosides was further tested on long-term erythroblast cultures. Medium was refreshed daily, reseeding cells at 1×10^6^ cells/mL. All data are displayed as mean ± SD (error bars; *n* = 3 donors, except for 0 h extracellular data of panel B, for which only 1 sample was measured). One-tailed paired Student’s t-test was used for the comparisons of panel C (𝐻_0_: 𝐶_−𝑛,𝑑𝑛_ = 𝐶_+𝑛,𝑑𝑛_, 𝐻_1_: 𝐶_−𝑛,𝑑𝑛_ > 𝐶_+𝑛,𝑑𝑛_; ns for non-significant, * for p < 0.05, ** for p < 0.01).

When nucleosides/deoxynucleosides were not added to the medium, hypoxanthine did not increase, as expected (Figure 6B; supplementary data 2). Allantoate, however, still increased to 70% of the levels observed in presence of additional nucleosides. This implies that allantoate, as a product of urate oxidation, is a metabolic byproduct of proliferating erythroblasts, possibly due to a high RNA turnover. The intracellular levels of ATP, ADP, and AMP appeared independent of the addition of nucleosides (Figure 6B). To evaluate the long-term effect of nucleosides removal from the medium on cell growth, erythroblasts were cultured with or without nucleosides/deoxynucleosides supplementation, with daily addition of fresh medium. Increased cell proliferation was observed under medium without exogenous nucleosides in the first 5 days of culture (Figure 6C).

### Oxidative stress as a limiting factor for cell growth

The accumulation of allantoate through urate oxidation suggests an active purine-mediated pathway of reactive oxygen species (ROS) generation in these cells. A state of oxidative stress in the cells may also be caused by the culture at atmospheric oxygen levels (hyperoxic, compared to *in vivo* conditions), or due to the high metabolic activity required to support rapid erythroblast proliferation. Thus, the activity of antioxidant pathways is likely an important parameter in erythroblast cultures. Specifically, control of oxidative stress by ascorbic acid and the glutathione pathway is crucial in erythropoiesis ^45^. Intracellular metabolomics showed depletion of the glutathione pool (GSH, GSSG, Figure 7A) along with ascorbate. Oxidized counterparts, dehydroascorbate, increased at 12 h only in high density cultures but later decreased well below starting levels (Figure 7). Glutathione or γ-l-glutamyl-l-cysteinyl glycine is synthesized from the γ-glutamyl cycle by γ-glutamyl-cysteine ligase and glutathione synthetase from glycine, glutamate, and cysteine. While glutamate accumulated extracellularly over the whole experiment, the other precursor glycine and the intermediate γ-glutamyl-cysteine first increased intracellularly until 24 h, followed by a decline well below starting levels. (Figure 7B).

**Figure 7.**
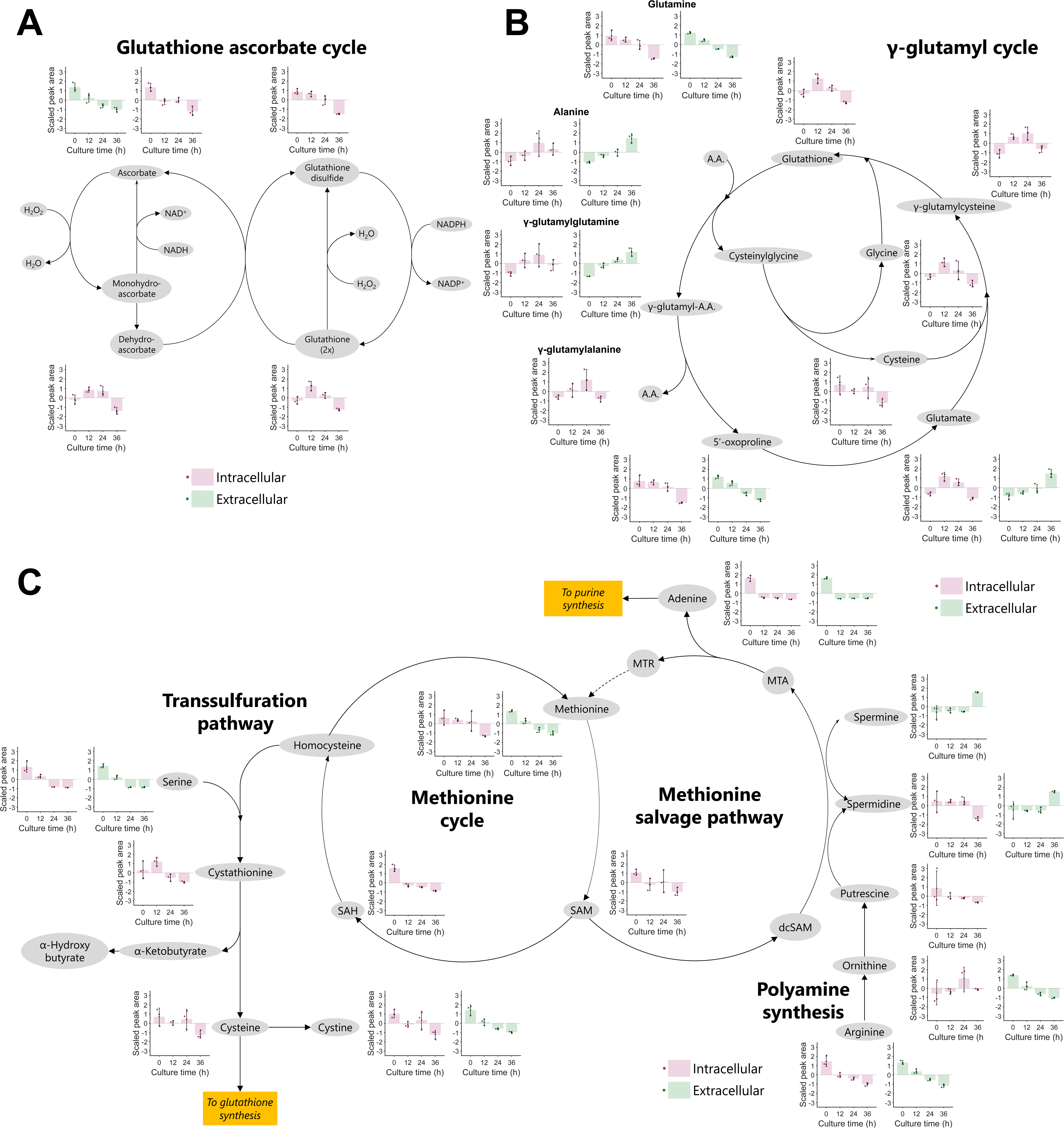
Depletion of glutathione and glutathione synthesis intermediates during erythroblast expansion cultures. Extracellular (green) and intracellular (red) levels of the glutathione-ascorbate cycle **(A)**, gamma-glutamyl cycle **(B)**, and methionine cycle + salvage pathway **(C)** for erythroblast (day 11) cultured at a starting cell concentration of 2.0×10^6^ cells/mL for 36 h in fresh proliferation medium. Some metabolites were only detected intra- or extracellularly. All data are displayed as mean ± SD (error bars; *n* = 3 donors).

Cysteine is considered to be the limiting substrate of *de novo* GSH synthesis ^46,47^. Cysteine can be supplemented to the media, but can also be synthesized from methionine via transsulfuration pathway. However, methionine is also decreased with time as early as 12h (Figure 7). All measured metabolites from both the transsulfuration pathway and the methionine cycle displayed a continuous decrease in concentration during 36 h. Furthermore, byproducts of the methionine salvage pathway accumulated. Particularly, the polyamines spermine and spermidine, accumulated extracellularly and intracelluarly (Figure 6 and 7). With these results, we hypothesize that exhaustion of erythroblast culture medium impacts the control of oxidative stress levels intracellularly through a decrease in glutathione pools and glutathione biosynthesis capacity.

## Discussion

Economically feasible production of cRBC for transfusion purposes requires efficient erythroblast expansion and differentiation, significant media cost reduction, and large-scale bioreactor cultivation ^4,34^. Cell density limitations in erythroblast expansion represent a clear challenge for the scale up of cRBC production. At a maximum erythroblast cell density of 2x10^6^ cells/mL, the production of a single transfusion unit (2x10^12^ cells) would require a 1000 L culture. The current study shows that exhaustion of small molecules (<3 kDa) rather than the accumulation of toxic components or the depletion of growth factors limits cell density during the expansion phase of erythroblast cultures. Indeed, SCF was supplied at 100 ng/mL (5.4 nM), well above reported c-Kit binding affinity (∼0.3 nM ^48^), such that receptor occupancy is expected to remain near-saturation despite substantial ligand depletion. In contrast, EPO was supplied at 1 IU/mL (0.36 nM), corresponding to sub-nanomolar concentrations close to the reported EPOR Kd (∼0.1-0.6 nM). Consequently, EPO signaling may be more sensitive to ligand depletion over time. However, supplementation of Epo or SCF did not rescue proliferation, suggesting that these growth factors were not limiting.

Here, untargeted metabolomics provided a lead to metabolic pathways affected by medium exhaustion. Supplementation of spent media (generated by harvesting media from high density cultures after 16 h) with fresh medium suggested that lack of nutrients likely caused the observed growth inhibition of erythroblast cultures. Nevertheless, supplementation of different media nutrients did not lead to a significant recovery in growth. Bayley et al observed similar results for umbilical cord-derived erythroid cultures supplemented with extra glucose, amino acids, and vitamins ^15^. It is possible that the observed erythroblast growth limitations are caused by the combined effect of multiple simultaneous nutrient limitations, as observed in other bioprocesses and exemplified by the rescue of erythroblast proliferation by full media supplementation ^49^.

Thus, feeding strategies must be designed to replenish depleted nutrients, and to remove or reduce the production of toxic metabolites that can become growth inhibiting at higher cell densities. We showed that continuous perfusion with fresh medium allows to overcome the observed cell density limitations in erythroblast cultures, albeit leading to large fresh medium requirements. Of note, we observed lower cell concentrations in the perfusion run compared to the predictions of the kinetic model ^33^. This may be explained by the different culture conditions used by Glen et al. which lacked glucocorticoids and thus allowing for fast cell divisions typical of terminal differentiation, opposite to our experimental model, which maintains cells in a transient renewal phase.

Determination of an optimal perfusion profile for erythroblast cultures can be determined experimentally, but our results indicate that *in silico* methods with growth models fitted to experimental data prior to experimental upscaling and validation may be more efficient. Such modelling will be sensitive to the hypothesized inhibition production mechanism. As an alternative to the kinetic model proposed by Glen et al., we also evaluated a simplified model, with growth-dependent inhibition and no inhibitor degradation fitted to the same cell concentration experimental data, for the determination of the optimal feeding profiles (model details available in Supplementary Methods). Although similar results were obtained using the kinetic model proposed by Glen et al. and this simplified model, significantly more medium is required for a situation in which there is no inhibition decay (which is a growth-inhibitory factor that decreases over time independently of dilution). As it is difficult to assert whether this mechanism takes place without knowing the set of metabolites that is being depleted during culture (for which we made a start in this work), the kinetic model without inhibition decay appears to fit better limitations in cell productivity observed in batch cultures.

We stress that a presumably complex dynamics of sets of growth inhibitory metabolites, decaying or not, over time requires more complex modeling and validation.

Erythroblast proliferation cultures require massive expansion levels to produce the number of RBCs of a single transfusion unit, and consequently, high uptake rates of amino acids to support the protein synthesis for new biomass production. We confirmed a generalized decrease of the amino acid pool in the medium during erythroblast proliferation, with glutamine having the largest consumption rate. Glutamine, often the most abundant amino acid in cell culture medium, is quickly consumed and used for synthesis of TCA cycle intermediates, nucleotides, and non-essential amino acids ^50,51^. Glutamine availability has also been shown to be critical for the commitment of hematopoietic stem cells to the erythroid lineage, being mainly used as direct precursor for nucleotide synthesis ^52^.

Serine was heavily consumed from the media, and intracellular levels were also low, similarly to what was previously observed in cultures of an immortalized erythroblast cell line ^53^. Serine is non-essential amino acid produced from the glycolysis pathway. Subsequently, serine can be converted to glycine by serine hydroxymethyltransferase (SHMT1/2). Glycin supplies onecarbon units required for nucleotide biosynthesis and the folate–methionine cycles, consistent with the observed decrease in methionine cycle intermediates. The extra carbon from converting serine to glycine may be routed to tetrahydrofolate forming 5,10-methylene-tetrahydrofolate, which is used in the synthesis of purines to support proliferation^54,55^. In addition, a total of 8 Gly molecules are required for a single heme unit, or 1.2 pg Gly per single fully hemoglobinized RBC containing ∼300 million Hb molecules. Low serine levels suggest that serine may become limiting in these pathways but without kinetic experiments it may also imply very efficient conversion to glycine. Well reflected in our data, a spike in glycine was observed at 12 and 24 h timepoints intracellularly.

Alanine and glutamate were the only amino acids that displayed extracellular accumulation during erythroblast proliferation. Alanine can be produced as an alternative to lactate from pyruvate via transamination, and its accumulation can inhibit the activity of pyruvate kinase, restricting glycolytic products from entering the TCA cycle ^56^. It has been hypothesized that alanine production via the activity of alanine aminotransferase (ALT; Pyr + Glu → Ala + αKG) coupled with the regeneration of glutamate via glutamate dehydrogenase (GDH; αKG + NH4+ → Glu) may be a resource to capture excess ammonia produced during culture ^57,58^.

Conversion of glutamine to the TCA cycle intermediate 2-oxoglutarate via glutamate is the main source of ammonia production ^21^. We have recently reported production rates of ammonia of 0.1 pmol/cell/day in erythroblast bioreactor and dish cultures, corresponding to ∼30% of the total glutamine uptake rates reported here ^26^. Although ammonia levels were <0.6 mM, below typical growth-inhibiting concentrations for other cells, higher concentrations are expected in cultures at higher cell densities. We suggest media optimization to be done by balancing amino acid concentrations according to the uptake rates reported in this work. Following this, an optimal perfusion rate sufficient to supply enough amino acids for erythroblast proliferation could be further implemented. This perfusion profile should also ensure the removal of toxic metabolites, while avoiding the excessive accumulation of metabolites that can be precursors to those metabolic byproducts. This could allow for the reduction of total medium requirements and the overall costs of the process.

Metabolite set enrichment analysis (MSEA) and pathway enrichment analysis (PEA) incorporate biological-relevant information (set of metabolites involved in the same pathway) missing in purely statistical methods such as PCA or clustering analysis. Following this approach, we could identify increased activity in the purine degradation pathway, causing transient accumulation of hypoxanthine. The expression of erythroid adenosine deaminase (eADA) is well documented and its levels are used as a diagnostic criterium in inherited anemia^59^. It is partly expressed as an ecto-enzyme to regulate the activation of adenosine receptors^60^. Addition of adenosine to culture medium may directly act on adenosine receptors and result in growth inhibitory signaling. In addition, adenosine deamination and hypoxanthine accumulation promotes ROS related injuries ^61^. Reducing nucleoside supplementation lowers both growth inhibitory pathways thereby promoting cell growth. This observation is in concordance with previous studies^62^ demonstrating that nucleoside supplementation in *in vitro* cultures enhances cell differentiation while reducing proliferation. It underscores the importance of metabolic modulation in improving culture conditions. Of note, the accumulation of allantoate was observed even when nucleosides were not supplemented in the medium, implying that allantoate is naturally produced in proliferating erythroblasts, possibly due to a high flux of the purine salvage pathway in cells that extensively change their transcriptome.

Oxidative stress imbalances were prominent in the erythroblast culture. Intracellular levels of GSH and its precursors are notorious for declining over time in erythroblast culture ^63^. Furthermore, levels of ascorbate, a potent antioxidant present in culture media ^45^, decreased concomitant with an increase in its oxidized form (dehydroascorbate). Nevertheless, accumulation of intracellular polyamines was observed over time in the culture. An increase in polyamine synthesis creates a low methylation, high redox buffer state. Polyamines protect against oxidative stress not primarily by classical redox chemistry, but by electrostatic shielding of nucleic acids, suppression of mitochondrial ROS generation, metal-mediated radical inhibition, and induction of stress adaptive programs such as autophagy ^64^. Consequently, this results in a shift in SAM metabolic flux from methylation toward cytoprotective polyamine synthesis. This result suggests that erythroblasts use polyamine biosynthesis as a compensatory mechanism to cope with limited levels of redox cofactors.

Despite our best efforts, some analytical constraints are linked to the present study. It is important to note that MSEA and PEA are strongly influenced by the selection of pathway and metabolite set databases, as these libraries differ in their coverage of reactions in the human metabolic network. However, the choice of analytical platform foremost defines the pool of identifiable metabolites ^41^. We used a rapid UHPLC-MS method that covers the central carbon and nitrogen metabolism ^65,66^. However, vitamins and heme metabolism intermediates, which are especially relevant in erythropoiesis, are generally poorly detected ^67^. To fill these gaps, assays targeting these pathways could be performed to further complement our metabolomics data.

Ultimately, the results reported here can be used to develop erythroblast culture medium and feeding profiles more rationally, considering the dynamic metabolic requirements of cultured erythroblasts. Currently, we are extending the approach shown in this study to the analysis of erythroid differentiation, in which drastic morphological changes take place, including organelle clearance and changes in membrane structural organization.

## Materials and Methods

### Erythroblast cell culture

Blood from healthy volunteer donors was collected and peripheral blood mononuclear cells (PBMCs) were purified by density centrifugation using Ficoll-Paque (density = 1.077 g/mL; 600g, 30 minutes; GE Healthcare; USA). Informed written consent was given by donors to give approval for research purposes, and was checked by Sanquin’s NVT Committee (approval file number NVT0258; 2012) in accordance with the Declaration of Helsinki and the Sanquin Ethical Advisory Council. Erythroblast were cultured from PBMCs as previously described ^68^, with minor modifications in the expansion medium composition: trace elements were omitted; cholesterol, oleic acid, and L-α-phosphatidylcholine were replaced by a defined lipid mix (1:1000; Sigma-Aldrich cat#L0288; USA). In short, PBMCs were cultured in the presence of human stem cell factor (hSCF; 100 ng/mL, produced in-house with HEK293T cells), erythropoietin (Epo; 2 IU/mL; EPREX®; Janssen-Cilag; Netherlands), dexamethasone (Dex; 1 µmol/L; Sigma-Aldrich), and interleukin-3 (IL-3; 1 ng/mL, first day only; Stemcell Technologies; Canada). After 7 days, a pro-erythroblast population (CD235alow/CD71^+^/CD49^+^) was obtained, and maintained in culture at a cell concentration of 0.7-2×106 cells/mL by daily feeding with fresh expansion medium prior to culture experiments. For cultures without nucleosides, medium without adenosine, guanosine, thymidine, cytidine, uracil, 2’-deoxyadenosine, 2’-deoxicytidine, and 2’-deoxyguanosine was used. Cell concentration was determined using an electric current exclusion with a size cutoff of 7.5 µm (CASY Model TCC, OLS OMNI Life Science, Germany; or Z2 Coulter Counter, Beckman Coulter, USA). Viability was determined using a hemocytometer and a dye exclusion method (Trypan Blue; Sigma).

### Bioreactor perfusion culture

The perfusion bioreactor culture was performed in a 500 mL glass bioreactor (MiniBio; Gettinge Biotechnology; Netherlands) with a working volume of 250 mL. A day 9 cell culture from PBMCs were seeded in the bioreactor at an initial cell concentration of 1×10^6^ cells/mL. Agitation was performed using a down-pumping marine impeller (100 rpm; diameter = 2.8 cm). Dissolved oxygen concentration was maintained at 2.9 mg/L (40% of the oxygen saturation concentration in water at equilibrium with air at 1 atm and 37°C) by sparging with air, and pH was controlled at 7.4 by sparging with CO_2_. Cell retention in the bioreactor was performed using an acoustic settler (BioSep 1 L; Applikon Biotechnology) run without recirculation and using a cyclic backflushing mode, as described before ^69,70^, set to the following parameters: ON time = 5-15 min, OFF time = 10-15 s, power = 3 W. The volumetric perfusion rate was adjusted daily based on the cell counting data to obtain an average cell-specific perfusion rate (CSPR) of 500 pL/cell/day, equivalent to the fresh medium utilization rate in the repeated batch cultivation method with half medium refreshment every 24h.

### Growth factor quantification

For quantification of extracellular Epo and hSCF concentrations, erythroblasts tissue static cultures were performed in proliferation medium with 2 IU/mL Epo (EPREX®; Janssen-Cilag; Netherlands) and 100 ng/mL hSCF (BioLegend cat#573908; USA). Supernatant samples were snap frozen, and stored at −80°C until measurement. After thawing on ice, samples were analyzed via sandwich enzyme-linked immunosorbent assays (ELISA; ab211647 and ab176109, Abcam; USA).

The average q-rates for amino acids and growth factors were calculated with Equation 1, where 𝑞_𝑖_ is the cell-specific consumption (>0) or production (<0) rate for the component 𝑖, 𝐶_𝑖,𝑡_ and 𝐶_𝑖,𝑡+1_ are the measured concentrations of component 𝑖 at the evaluated timepoints (𝑡 and 𝑡 + 1), and 𝐶_𝑋,𝑡_ and 𝐶_𝑋,𝑡+1_ are the measured cell concentrations at the same timepoints. The average growth rate 𝜇 in that same time interval was calculated using Equation 2.

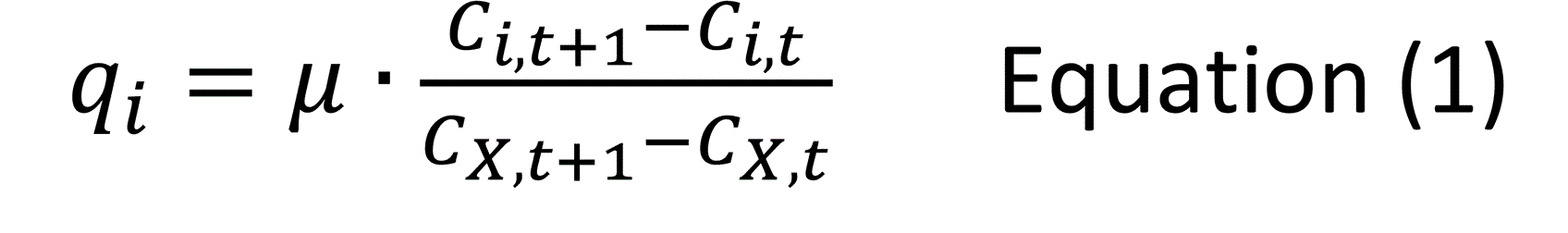

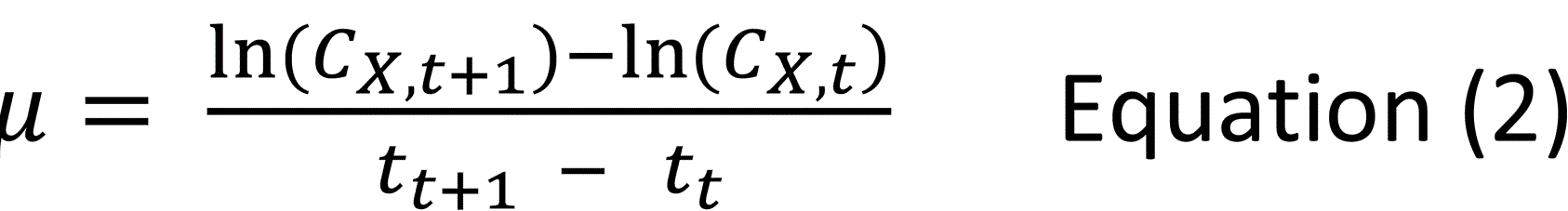

### Spent medium analysis and fractionation

Erythroblasts were expanded from PBMCs for 11 days and subsequently seeded in fresh medium at cell concentrations between 1 and 15×10^6^ cells/mL, as indicated. Cells were cultured for 16 h, and spun down (600g, 5 min). Spent medium was filtrated (0.22 µm), snap frozen in liquid nitrogen, and stored at −80°C until further use. Upon thawing, day 9 erythroblasts were washed, resuspended in fresh medium, spent medium or a 1:1 (v/v) combination of both, and cultured for 48 h without medium refreshment. For medium fractionation experiments, fresh or spent medium was filtered using an Amicon Ultra-15 3kDa centrifugal filter, following manufacturer’s protocol (Millipore Merck; Germany), resulting in filtrate and retentate fractions. Day 9 erythroblasts were seeded in dishes in combinations of retentate and filtrate of fresh or spend medium, as indicated.

### Spent medium supplementation

The effect of spent medium supplementation with specific fractions of IMDM was evaluated using previously prepared solutions with respective IMDM components. For basal IMDM (bIMDM), a solution containing 1.49 mM CaCl_2_, 0.81 mM MgSO^4^, 4.43 mM KCl, 36 mM NaHCO_3_, 77.1 mM NaCl, 1.04 mM NaH_2_PO_4_, 0.75 µM KNO_3_, 25 mM HEPES, and 25 mM D-glucose was prepared with demineralized water. For selenite supplementation, Na_2_SeO_3_ was added to bIMDM to a final concentration of 0.1 µM (bIMDM+Se). To test the effect of amino acid supplementation, bIMDM was supplemented with essential (bIMDM+EAA) amino acids only, or with both essential and non-essential amino acids (bIMDM+EAA+NEAA). Amino acids in the prepared media were at 2X the concentration in commercial IMDM, so when combining 1:1 with spent medium any depleted amino acid would be restored to the levels originally present in IMDM. Final concentrations of essential amino acids in bIMDM+EAA and bIMDM+EAA+NEAA were 1.6 mM L-lysine, 0.93 mM L-tyrosine, 0.58 mM L-cystine, 0.8 mM L-arginine, 0.4 mM L-histidine, 1.6 mM L-isoleucine, 0.4 mM L-methionine, 0.8 mM L-phenylalanine, 1.6 mM L-leucine, 1.6 mM L-threonine, 0.16 mM L-tryptophan, 1.6 mM L-valine, and 4.0 mM L-glutamine. For non-essential amino acids, final concentrations in bIMDM+EAA bIMDM+EAA were 0.8 mM L-serine, 0.45 mM L-aspartate, 0.69 mM L-proline, 0.8 mM glycine, 0.38 mM L-asparagine, 0.56 mM L-alanine, and 1.0 mM L-glutamate. Similarly, bIMDM was supplemented with vitamins (bIMDM+Vit) to have final concentrations 2X higher relative to commercial IMDM. For this, MEM vitamin mix (ThermoFisher cat#11120052), biotin and vitamin B12 were added to bIMDM to final concentrations as follows: 8 mg/L nicotinamide, 8 mg/L cholin chloride, 8 mg/L Ca-D-pantothenate, 8 mg/L pyridoxal HCl, 8 mg/L thiamine HCl, 0.8 mg/L riboflavin, 8 mg/L folic acid, 16 mg/L myo-inositol, 26 µg/L biotin, and 26 µg/L vitamin B12. Epo, hSCF and dex were added to the final media combinations (e.g. 1:1 spent medium and bIMDM) to have final concentrations of 2 U/mL, 100 ng/mL and 1 µM, respectively.

### Targeted profiling of culture supernatant

Measurement of amino acid concentrations in supernatant samples was determined via gas chromatography-mass spectrometry (GC-MS) and isotope dilution mass spectrometry (IDMS), as previously described ^71^. Briefly, culture samples were spun down (600g, 5min), and the supernatant was freeze dried, reconstituted and derivatized using acetonitrile and N-tert-Butyldimethylsilyl-N-methyltrifluoroacetamide (MTBSTFA), and subsequently analyzed by GC-MS. IDMS for amino acid quantification was performed using a U-^13^C-labelled cell extract as an internal standard mix.

### Extra- and intracellular untargeted UHPLC-MS metabolomics

Day 11 erythroblasts of three independent donors were washed twice with phosphate-buffered saline (PBS), resuspended in fresh medium, and cultured for 36 h under static conditions without medium refreshment. Samples containing 3×10^6^ cells were spun down (600g, 5 min, 4°C), and both cell pellet and supernatant were snap frozen in liquid nitrogen and stored at −80°C until extraction. Metabolite extraction was performed using an ice-cold methanol:acetonitrile:water buffer (5:3:2) by vortexing for 30 minutes at 4°C. Analyses were performed using a Vanquish UHPLC system (Thermo Fischer Scientific; USA) coupled online to a Q Exactive mass spectrometer, as previously described ^65^. Metabolite assignment was performed using MAVEN ^72^. Data cleaning was performed in three stages: (i) pre-filtering of metabolites that did not show significant differences in peak areas (Student t-test, p>0.05) at any condition or timepoint compared to blank samples, (ii) missing value imputation (missing values replaced by 1/5 of the smallest positive value in the original dataset), and (iii) filtering based on the quality control (QC) samples (if a metabolite showed a relative standard deviation > 25% in the QC samples, it was not considered for further analysis).

### Data analysis and statistics

For analysis of metabolomics data, principal component analysis (PCA) was performed using the peak area data for each annotated metabolite after glog transformation and mean-centering. k-means clustering was performed using the ComplexHeatmap R package ^73^, using the clustree and factoextra packages to determine the optimal number of clusters ^74^. For functional analysis, metabolites were first mapped to the Human Metabolome Database (HMDB). Metabolite-set quantitative enrichment analysis (MSQEA) and pathway quantitative enrichment analysis (PQEA) of the complete set of metabolites and of individual k-means clusters were performed using MetaboAnalystR v3 (package version 3.0.3 amap_0.8-18) ^75,76^, and a metabolite set library derived from the Small Molecule Pathway Database ^77,78^. All R code used to process and analyze the metabolomics data is available at request. Significance levels shown in barplots were calculated using one or two-tailed paired Student’s t-test (ns for non-significant, * for *p* < 0.05, ** for *p* < 0.01, *** for *p* < 0.005).

### Mathematical modelling of growth inhibition

Erythroblast growth inhibition was modelled using kinetic unstructured unsegregated mathematical models described with a set of three ordinary differential equations (cell concentration, and non-dimensionalized inhibitor concentration). Detailed equations for the growth models and for the simulations of bioreactor cultures under different feeding regimes are available in Supplementary Methods.

## Author Contributions

Conceptualization, JSGM, MvL, AW, VPG, and EvdA; methodology, JSGM, NY, MvL, and AW; software, JSGM and IvL; formal analysis, JSGM, IvL, JR, and AD; investigation, JSGM, NY, and JR; resources, JR, AD; data curation, JSGM; writing—original draft preparation, JSGM, MvL, and AW; writing review and editing, VPG, EvdA, LvdW, AD; visualization, JSGM, MvL, and AW.; supervision, MvL, AW, EvdA, and AD; project administration, MvL; funding acquisition, MvL, EvdA, and AW. All authors have read and agreed to the published version of the manuscript.

## Institutional Review Board Statement

The study was conducted in accordance with the Declaration of Helsinki, and approved by Sanquin’s NVT Committee (approval file number NVT0258; 2012) and the Sanquin Ethical Advisory Board. Informed Consent Statement: Informed written consent was obtained from all donors to give approval for the use of waste material (including buffy coats used for the isolation of PBMCs) for research purposes.

## Data Availability Statement

The processed metabolomics data presented in this study are included with the submission as .xlsx files (Supplementary data1 and Supplementary data2). Raw files will be available upon request.

## Acknowledgments

We thank Arjan Hoogendijk and John van Dam (Bioinformatics, Sanquin Research) for technical help and advice on the analysis and visualization of metabolomics data. We also thank Patricia van Dam (Department of Biotechnology, Delft University of Technology) for the technical support in the determination of the amino acid profiles of the supernatant samples.

## Figure legends

