## supplemental data for "Small-molecule consumption drives metabolic stress and restricts erythroblast expansion in high-density cultures"

**Supplementary Methods**

**Kinetic models for erythroblast growth**

The kinetic growth model proposed by Glen et al. [1,2] was considered for the numerical simulations of erythroblast cultures under different feeding regimes. Briefly, this model assumes that erythroblasts (cell concentration = $C_X$) can proliferate under exponential growth with a maximum growth rate $\mu_{max}$. Accumulation of an inhibitory factor (non-dimensionalized concentration = $C_I^*$; arbitrary concentration units), which can be interpreted as either the accumulation of a toxic metabolite or the depletion of a growth-limiting metabolite, leads to a decrease in the growth rate. The dose-response curve for this inhibition is described by a sigmoidal function with two parameters: the inhibition threshold $k_c^*$ and the inhibitor sensitivity $k_s^*$ (Equation A1).

$$\mu = \mu_{max} \cdot (1 + \exp(-k_s^*(k_c^* - C_I^*))^{-1} \qquad \text{Equation (A1)}$$

The inhibitor $I$ is produced in a growth-dependent manner. An inhibition decay term (first order kinetics) is included, with an inhibitor decay rate $r_d$. Equations A2 and A3 describe the concentration of cells and inhibitory factor for a batch culture.

$$\frac{d\,C_X}{dt} = \mu\, C_X \qquad \text{Equation (A2)}$$

$$\frac{d\,C_I^*}{dt} = \frac{d\,C_X}{dt} - r_d C_I^* = \mu C_X - r_d C_I^* \qquad \text{Equation (A3)}$$

After fitting of this model to experimental data from erythroblast cultures under different treatments (batch growth; reset of cell concentration by removing all cells from a batch culture and inoculating the spent medium with fresh proliferating cells; medium dilution by harvesting cells and reinoculation in a defined ratio of spent and fresh medium), Glen et al. obtained the following best fit parameters: $\mu_{max} = 0.0541$ 1/h, $k_c^* = 3.4$ 1/h, $k_s^* = 5.6$ 1/h, $r_d = 0.014$ 1/h.

A simplified kinetic model was developed for this study in which: (1) the inhibitory factor is produced in a growth-dependent manner, (2) the inhibitor affects growth following a Hill dose-response curve, and (3) no inhibition decay is considered. Equation A4 describes the effect of the inhibitor in cell growth. The parameters of this model are the maximum growth rate $\mu_{max}$, the half-saturation concentration $K_I$, the Hill coefficient $n_I$, and the yield of inhibitor on cells $Y_{I/X}$. Equations A5 and A6 describe the concentrations of cells and inhibitory factor in a batch culture.

$$\mu = \mu_{max} \cdot \left(1 - \frac{1}{1+(K_I/C_I)^{n_I}}\right) \qquad \text{Equation (A4)}$$

$$\frac{d\,C_X}{dt} = \mu\, C_X \qquad \text{Equation (A5)}$$

$$\frac{d\,C_I}{dt} = Y_{I/X}\,\frac{d\,C_X}{dt} = Y_{I/X}\,\mu\, C_X \qquad \text{Equation (A6)}$$

In a similar manner as in Glen et al.' approach, the concentration of the inhibitory factor can be non-dimensionalized using the half-saturation constant ($C_I^* = C_I/K_I$), reducing the number of parameters of the model. A new parameter $\alpha_I^* = Y_{I/X}/K_I$ can be defined, resulting in Equations A7 and A8 describing the growth inhibition dose-response curve and the inhibitor concentration over time in a batch process, respectively:

$$\mu = \mu_{max} \cdot \left(1 - \frac{1}{1+\left(1/C_I^*\right)^{n_I}}\right) \qquad \text{Equation (A7)}$$

$$\frac{d\,C_I^*}{dt} = \alpha_I^*\,\mu\, C_X \qquad \text{Equation (A8)}$$

This model was fitted to the experimental dataset obtained by Glen et al. [1]. The fitting was performed minimizing the total sum of squared relative errors of cell concentrations for all treatments reported by Glen et al., weighted by the number of measurements of each treatment. A global optimization strategy was used as initial step (genetic algorithm), followed by local optimization using the Nelder-Mead simplex algorithm (MATLAB ver. R2018b; Mathworks; USA). Fitted parameters were: $\mu_{max} = 0.04$ 1/h, $\alpha_I^* = 0.32 \times 10^{-6}$ mL/cell, $n_I = 9.5$. Note that the non-dimensionalized inhibitor concentrations in both models are not equivalent, as the non-dimensionalization is performed using parameters from different dose-response inhibition models.

**Numerical simulations of erythroblast bioreactor cultures**

Bioreactor cultures under different feeding regimes were simulated solving numerically the differential equations describing the concentration of cells and (non-dimensionalized) inhibitor factor. Equations A9 and A10 show the generic mass balance for cells and inhibitory factor that were used for all feeding regimes. Equation A11 describes the change in liquid volume in the reactor:

$$\frac{d\,C_X}{dt} = \frac{1}{V}\left(-F_{out}C_{X,out} + \mu\,C_X\,V - C_X\frac{dV}{dt}\right) \qquad \text{Equation (A9)}$$

$$\frac{d\,C_I^*}{dt} = \frac{1}{V}\left(-F_{out}C_{I,out}^* + \alpha_I^*\,\mu\,C_X\,V - C_I^*\frac{dV}{dt}\right) \qquad \text{Equation (A10)}$$

$$\frac{dV}{dt} = F_{in} - F_{out} \qquad \text{Equation (A11)}$$

Where $F_{in}$ and $F_{out}$ are the volumetric inflow and outflow rates, and $V$ is the working volume of the bioreactor at any specific timepoint. Cell growth rate is calculated at each timepoint using Equation A4 or A7, depending on the growth inhibition model being evaluated. The value of $\alpha_I^*$ is equal to 1 for Glen et al. model.

The inflow and outflow terms are defined depending on the feeding regime. For batch mode, $F_{in} = F_{out} = 0$, resulting in $\frac{dV}{dt} = 0$, and Equations A5 and A8. For both fed batch regimes (FBconst and FBexp), $F_{out} = 0$, and the feeding rate was normalized by the initial working volume of the reactor $D = F_{in}/V_0$, resulting in Equations A12, A13 and A14. For FBconst, the value of D is constant, while for FBexp it is described by $D = D_0\exp(\theta \cdot t)$.

$$\frac{d\,C_X}{dt} = \frac{1}{V}\left(\mu\,C_X\,V - C_X\frac{dV}{dt}\right) \qquad \text{Equation (A12)}$$

$$\frac{d\,C_I^*}{dt} = \frac{1}{V}\left(\alpha_I^*\,\mu\,C_X\,V - C_I^*\frac{dV}{dt}\right) \qquad \text{Equation (A13)}$$

$$\frac{dV}{dt} = D\,V_0 \qquad \text{Equation (A14)}$$

For both perfusion regimes (Pconst and PCSPR), it is assumed that the working volume in the reactor is constant ($F_{out} = F_{in}$), that no cells are lost in the permeate ($C_{X,out} = 0$), and that the reactor is well-mixed for the inhibitory factor ($C_{I,out}^* = C_I^*$). The perfusion rate is expressed normalized by the working volume of the reactor $P = F_{out}/V$, resulting in Equations (A15), (A16) and (A17). For Pconst, the value of P is constant, while for PCSPR it is a function of the cell-specific perfusion rate CSPR and the cell concentration in the reactor ($P = CSPR \cdot C_X$).

$$\frac{d\,C_X}{dt} = \frac{1}{V}\left(\mu\,C_X\,V - C_X\frac{dV}{dt}\right) \qquad \text{Equation (A15)}$$

$$\frac{d\,C_I^*}{dt} = \frac{1}{V}\left(-P\,V\,C_I^* + \alpha_I^*\,\mu\,C_X\,V - C_I^*\frac{dV}{dt}\right) \qquad \text{Equation (A16)}$$

$$\frac{dV}{dt} = 0 \qquad \text{Equation (A17)}$$

To be able to compare different feeding regimes, the feeding parameters of each feeding regime were optimized under the same constraints: production of $30\times10^9$ cells in 4 days in a 1 L bioreactor inoculated at a starting cell concentration of $1\times10^6$ cells/mL in fresh medium free of the inhibitory factor. The objective function of the minimization problem was the total volume of medium required during the process.

**Supplementary Figures**

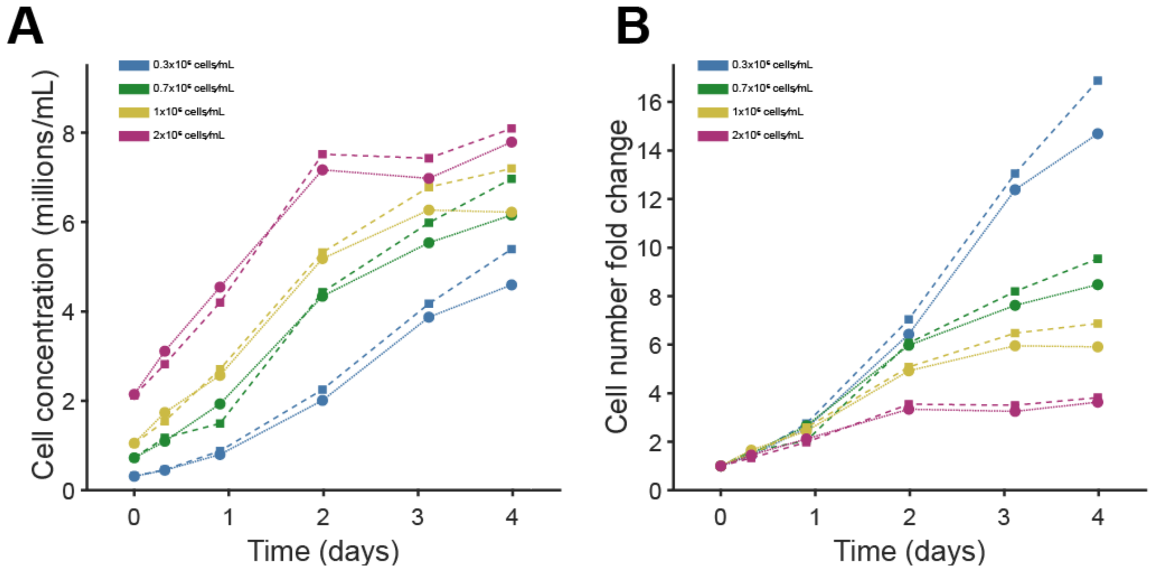


**Supplementary Figure S1. Growth in erythroblast cultures without medium refreshment.** PBMC-derived day 10 erythroblasts from two independent donors (donor 1: dashed line; donor 2: solid line) were inoculated in fresh proliferation medium at different starting cell densities (0.3, 0.7, 1 and $2\times10^6$ cells/mL; indicated by colors). Cell concentration **(A)** and cell number fold change relative to the start of the culture **(B)** are displayed.

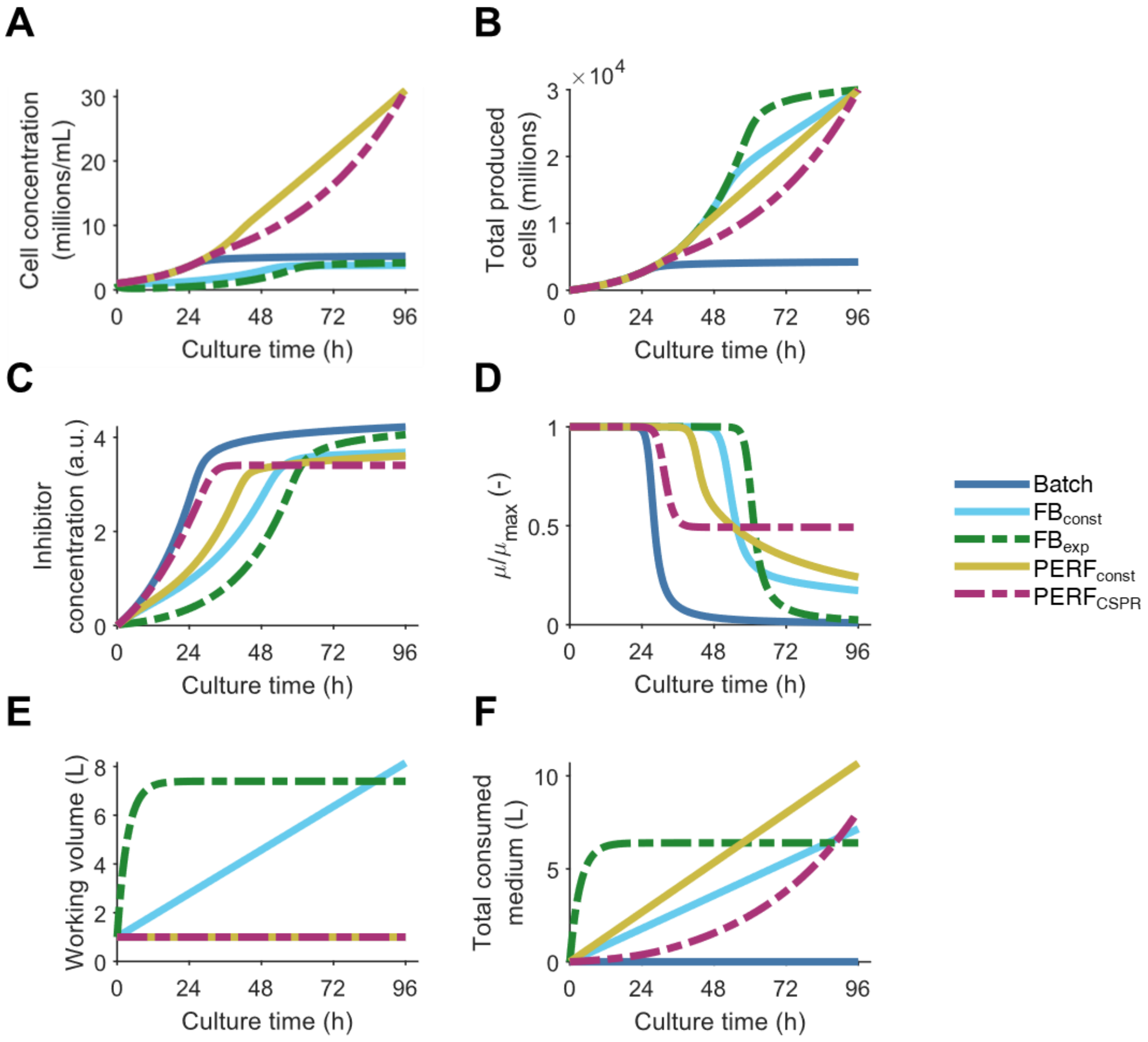


**Supplementary Figure S2. Evaluation of feeding strategies in erythroblast bioreactor cultures with alternative growth inhibition model.** Bioreactor cultures with different feeding strategies (batch, fed batch with constant feed rate $FB_{const}$, fed batch with exponentially increasing feed $FB_{exp}$, perfusion at a constant rate $PERF_{const}$, perfusion at a constant cell-specific perfusion rate $PERF_{CSPR}$) were simulated, using the kinetic growth model for erythroblast proliferation proposed by Glen et al.[1] Similar to the model used for Figure 2, this model includes the production of a putative inhibitor in a growth-dependent manner. However, the dose-response curve to this inhibitor is described using a sigmoidal equation. Furthermore, inhibition decay is also considered (first order kinetics). Except for batch mode, the feeding parameters of each feeding regime were optimized to achieve a production of $30\times10^9$ cells in 4 days in a 1 L bioreactor inoculated at a starting cell concentration of $1\times10^6$ cells/mL, minimizing the medium volume requirement. Cell concentration **(A)**, total produced cells **(B)**, concentration of the putative inhibitor (arbitrary units; **C**), magnitude of the growth inhibition calculated as the ratio of the instantaneous growth rate μ and the maximum growth rate μmax **(D)**, total volume of medium consumed **(E)**, and reactor working volume **(F)** are displayed for all evaluated feeding regimes. Equations used for the growth model and for the bioreactor simulations are available in Supplementary Methods. Optimized parameters for each feeding regime are included in Supplementary Table S1.

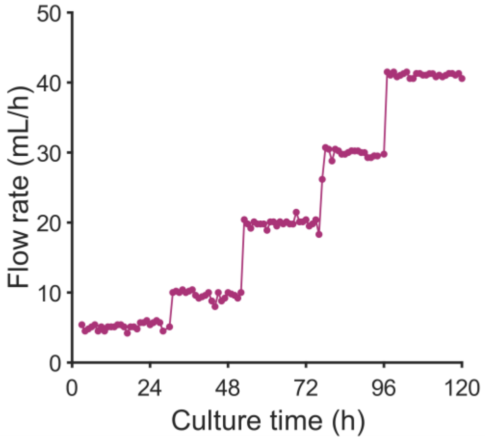


**Supplementary Figure S3. Perfusion rate in bioreactor with acoustic cell retention.** Erythroblasts were cultured for 5 days in a 0.5 L stirred tank bioreactor (working volume = 250 mL), using an Applikon BioSep for cell retention. The permeate flow rate was increased sequentially according to the daily measurements of cell concentration, targeting an average daily cell-specific perfusion rate (CSPR) of 500 pL/cell/day.

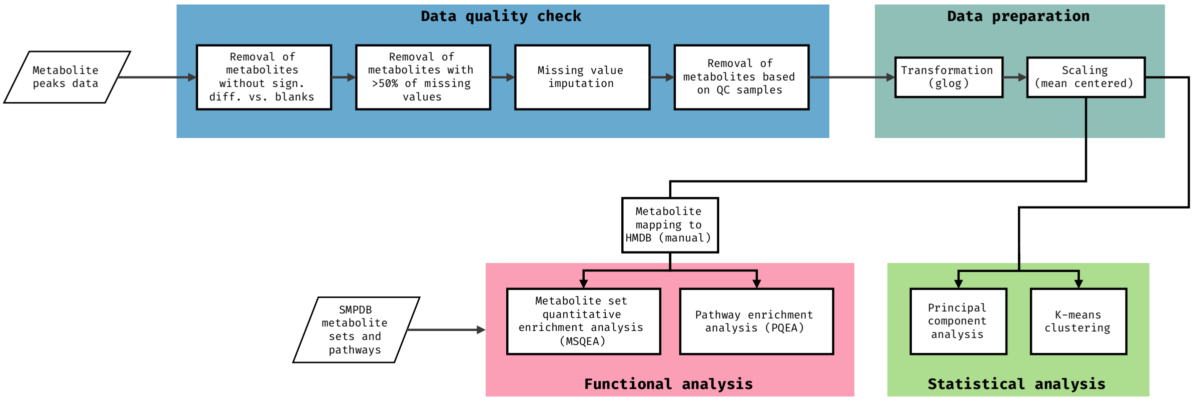


**Supplementary Figure S4. Workflow for analysis of untargeted metabolomic data.** Metabolite peak data obtained after acquisition (using a UHPLC-MS Vanquish coupled online to a Q Exactive mass spectrometer; Thermo Fischer Scientific; USA), validation, and annotation (using MAVEN), was used as input for the metabolomics analysis workflow. Data cleaning was performed in three stages: (i) pre-filtering of metabolites with peak areas not significantly different to blank samples (Student t-test; $p$<0.05), (ii) missing value imputation (missing values replaced by 1/5 of the smallest positive value in the original dataset), and (iii) filtering based on the quality control (QC) samples (if a metabolite showed a relative standard deviation > 25% in the QC samples, it is not considered for further analysis). Following this, peak data was transformed (glog method) and scaled (mean centered). Processed peak data was used for statistical analysis (principal component analysis and *k*-means clustering). For functional analysis, metabolites were first mapped to the Human Metabolome Database (HMDB). Metabolic set quantitative enrichment analysis (MSQEA) and pathway quantitative enrichment analysis (PQEA) was performed using MetaboAnalystR using a database of metabolite sets and pathways from the Small Molecule Pathway Database (SMPDB).

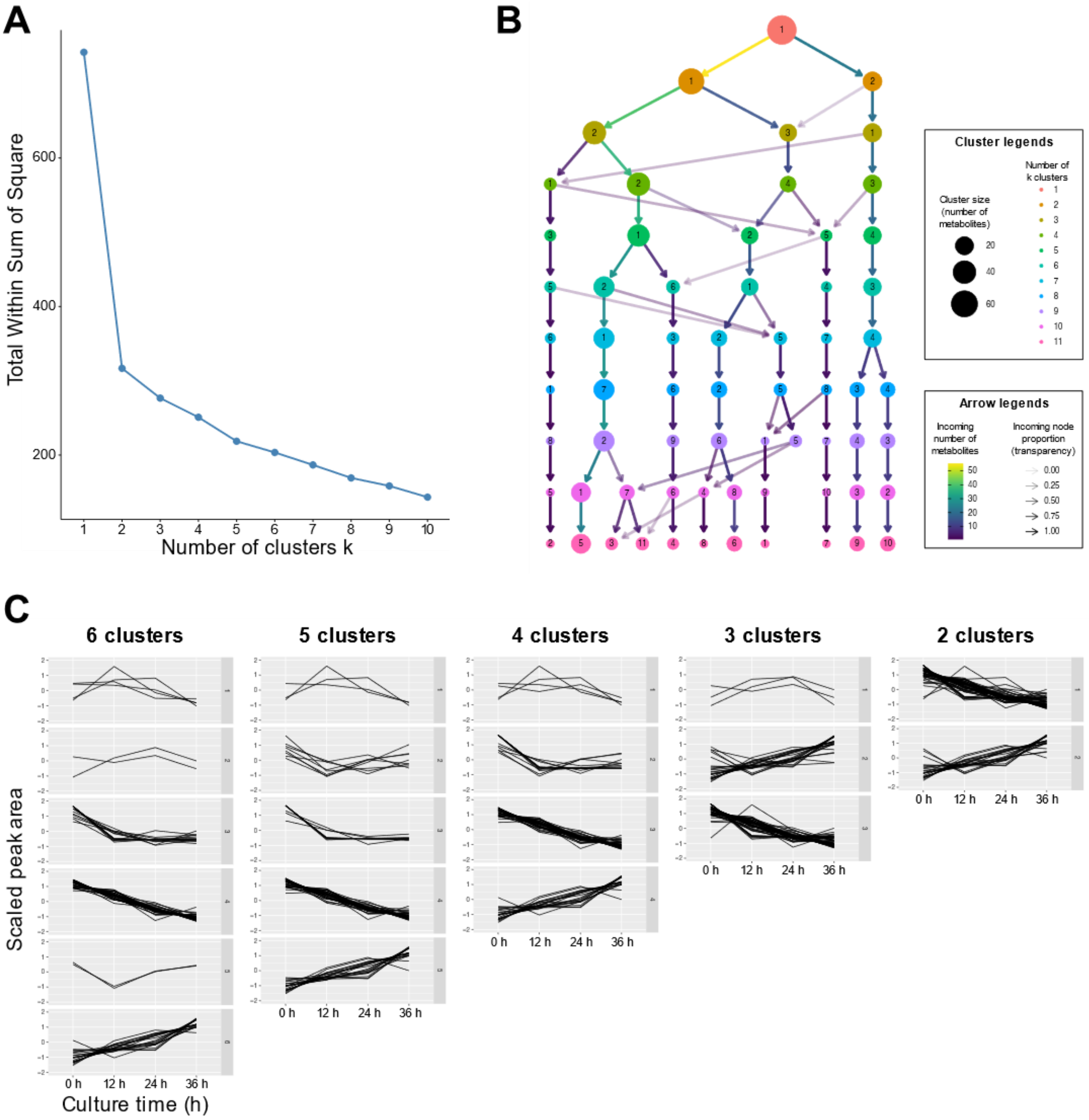


**Supplementary Figure S5. Determination of optimal number of *k*-clusters for extracellular metabolomics data of high cell concentration erythroblast cultures. (A)** Effect of number of *k*-clusters on the total explained variation. Increasing the number of clusters leads to a higher percentage of the total variation explained by the clusters, here measured as a decrease in the total sum of squares. **(B)** Effect of increasing cluster number on the distribution of metabolites in the *k*-clusters. Over-clustering of data can be seen as an increase in incoming arrows to a newly generated clusters, representing a reassignment of metabolites previously in other clusters in the new cluster. An example of this can be seen when going from 6 to 7 clusters, with cluster 5 receiving metabolites from three different clusters. **(C)** Trends of scaled peak area for each cluster as cluster number increases from 2 to 6. Results from all panels were generated using the function *fviz_nbclust* from the R package *factoextra*.

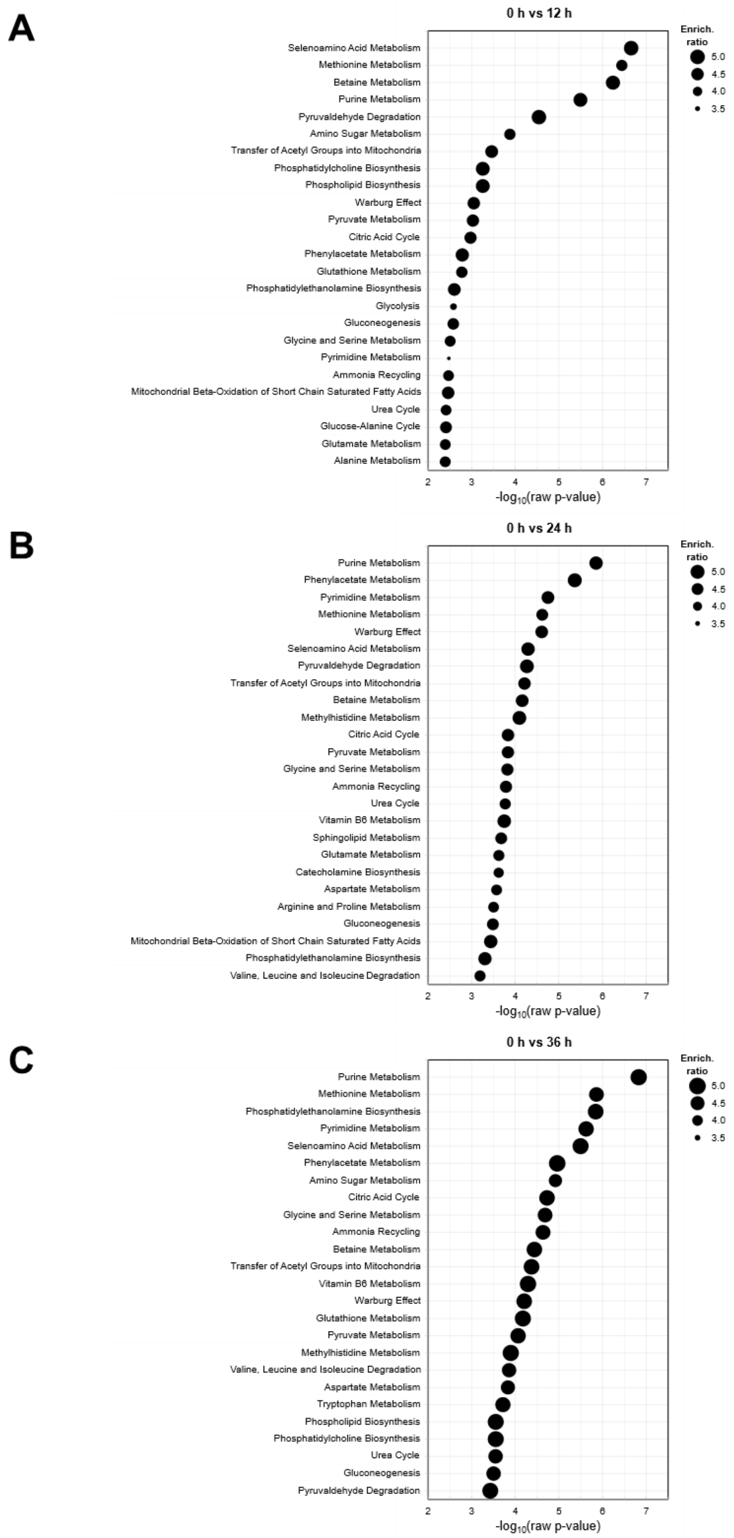


**Supplementary Figure S6. Metabolite set quantitative enrichment analysis (MSQEA) of extracellular metabolome data of proliferating erythroblast cultures.** After mapping the identified metabolites to the Human Metabolome Database (HMDB), metabolite-set quantitative enrichment analysis (MSQEA) was performed. Supernatant untargeted metabolomics data for cultures started at a high cell concentration ($2.0\times10^6$ cells/mL) was used for this analysis. Results for the pairwise comparisons of 12h **(A)**, 24h **(B)**, and 36h **(C)** relative to the start of the culture (0h) are displayed. MSQEA was performed using the Small Molecule Pathway Database (SMPDB) available in MetaboAnalystR v3. For each metabolite set or pathway, the enrichment level and *p*-value are displayed (x-axis and size of the circle, respectively). Only the top 25 most enriched metabolic pathways are displayed.

### Supplementary Tables

**Supplementary Table S1. Optimized operating parameters for evaluated feeding profiles.** Parameters for fed-batch and perfusion feeding strategies were optimized to achieve a production of 30×10$^{9}$ cells in 4 days in a 1 L bioreactor inoculated at a starting cell concentration of 1×10$^{6}$ cells/mL, minimizing total medium consumption. Optimized parameters were as follow: feeding rate normalized by initial culture volume ($F/V_0 = D$; 1/h) for the fed batch culture using constant feed rate $FB_{const}$; initial feeding rate normalized by initial culture volume ($F_0/V_0 = D_0$; 1/h) and exponential increase rate of the feeding rate ($\theta$; 1/h) for the fed batch culture with exponentially increasing feed $FB_{exp}$; perfusion rate normalized by culture volume ($F/V_0 = P$; 1/h) for the perfusion culture at a constant perfusion rate $PERF_{const}$; cell-specific perfusion rate (CSPR; pL/cell/day) for the perfusion at a constant CSPR $PERF_{CSPR}$. Two different growth inhibition models were used for the bioreactor simulations: the growth inhibition model proposed by Glen et al. including inhibition decay, and a simplified model without inhibition decay (This study) fitted to the same dataset. Equations used for the growth model and for the bioreactor simulations are available in Supplementary Methods.

| Feeding strategy | Optimized feeding parameters | This study | Glen et al. (2018) |
|---|---|---|---|
| $FB_{const}$ | D (1/h) | 0.086 | 0.075 |
| $FB_{exp}$ | $D_0$ (1/h) | 0.49 | 1.89 |
| | θ (1/h) | -0.064 | -0.30 |
| $PERF_{const}$ | P (1/h) | 0.19 | 0.11 |
| $PERF_{CSPR}$ | CSPR (pL/cell/day) | 297 | 187 |

**Supplementary Table S2. Quantification of amino acid concentrations in erythroblast proliferation cultures.** Erythroblast from two donors were cultured from PBMCs for 10 days, seeded in fresh medium at a starting cell density of 1.0×10$^{6}$ cells/mL, and cultured for 2 days. Supernatant amino acid concentrations at days 0 and 2 of culture were measured by GC-MS. Control samples in medium without cells were also measured. Data not available (n.a.) for arginine, as it could not be measured with the used method.

| | | Extracellular concentration (µmol/L) | | | | | | | | | | | | | | | | | | | |
|---|---|---|---|---|---|---|---|---|---|---|---|---|---|---|---|---|---|---|---|---|---|
| **Condition** | **Day** | **Ala** | **Arg** | **Asn** | **Asp** | **Cys** | **Gln** | **Glu** | **Gly** | **His** | **Ile** | **Leu** | **Lys** | **Met** | **Phe** | **Pro** | **Ser** | **Thr** | **Trp** | **Tyr** | **Val** |
| No cells | 0 | 296 | n.a. | 265 | 240 | 144 | 2454 | 547 | 436 | 264 | 797 | 764 | 890 | 181 | 395 | 339 | 366 | 841 | 91 | 467 | 825 |
| No cells | 2 | 303 | n.a. | 238 | 235 | 202 | 2129 | 563 | 397 | 280 | 835 | 784 | 909 | 186 | 412 | 362 | 395 | 930 | 84 | 479 | 828 |
| Donor 1 | 0 | 285 | n.a. | 239 | 235 | 137 | 2346 | 503 | 406 | 233 | 785 | 739 | 868 | 175 | 390 | 339 | 349 | 771 | 85 | 444 | 779 |
| Donor 1 | 2 | 520 | n.a. | 192 | 244 | 135 | 796 | 721 | 328 | 155 | 530 | 420 | 643 | 130 | 310 | 328 | 10 | 728 | 56 | 397 | 523 |
| Donor 2 | 0 | 303 | n.a. | 224 | 231 | 224 | 2410 | 529 | 417 | 247 | 806 | 770 | 910 | 175 | 399 | 352 | 364 | 806 | 91 | 452 | 812 |
| Donor 2 | 2 | 432 | n.a. | 172 | 254 | 111 | 752 | 679 | 292 | 145 | 547 | 429 | 646 | 129 | 306 | 316 | 17 | 718 | 61 | 392 | 529 |

**Supplementary Table S3. Cell-specific consumption and production rates of amino acids in erythroblast proliferation cultures.** Erythroblast from two donors were cultured from PBMCs for 10 days, seeded in fresh medium at a starting cell density of 1.0×10$^6$ cells/mL, and cultured for 2 days. Supernatant amino acid concentrations at days 0 and 2 of culture were measured by GC-MS (concentrations available in Supplementary Table S2). The cell specific consumption/production rate ($q_i$; <0 if consumed, >0 if produced) for each amino acid was determined using the average growth rate between day 0 and 2 (Equation 2), and the initial and final concentration of the amino acid in the supernatant (Equation 1). Data not available (n.a.) for arginine, as its concentration could not be quantified with the used method.

| | $q_i$ (nmol/million cells/day) | | | | | | | | | | | | | | | | | | | |
|---|---|---|---|---|---|---|---|---|---|---|---|---|---|---|---|---|---|---|---|---|
| **Donor** | **Ala** | **Arg** | **Asn** | **Asp** | **Cys** | **Gln** | **Glu** | **Gly** | **His** | **Ile** | **Leu** | **Lys** | **Met** | **Phe** | **Pro** | **Ser** | **Thr** | **Trp** | **Tyr** | **Val** |
| 1 | 43.0 | n.a. | -8.5 | 1.6 | -0.4 | -283 | 39.9 | -14.2 | -14.2 | -46.6 | -58.3 | -41.2 | -8.3 | -14.7 | -2.0 | -61.8 | -7.8 | -5.4 | -8.7 | -46.8 |
| 2 | 24.4 | n.a. | -10.0 | 4.2 | -21.3 | -314 | 28.4 | -23.8 | -19.4 | -49.2 | -64.7 | -50.0 | -8.7 | -17.6 | -6.8 | -65.9 | -16.8 | -5.7 | -11.4 | -53.6 |
| Average | 33.7 | n.a. | -9.2 | 2.9 | -10.9 | -299 | 34.1 | -19.0 | -16.8 | -47.9 | -61.5 | -45.6 | -8.5 | -16.1 | -4.4 | -63.8 | -12.3 | -5.5 | -10.1 | -50.2 |
